# Unequal requirement of KAI2 for AM symbiosis across angiosperms

**DOI:** 10.64898/2026.05.03.722480

**Authors:** Kees Buhrman, Salar Torabi, Samy Carbonnel, Kartikye Varshney, Philipp Chapman, Amit Fenn, Maxim Messerer, Götz Hensel, Nadia Kamal, Caroline Gutjahr

## Abstract

Development of arbuscular mycorrhiza (AM), a symbiosis between plants and beneficial *Glomeromycotan* fungi, is largely under plant control. Several genes, required for AM development, are proposed to be regulated by the karrikin signalling module, comprising the alpha/beta hydrolase receptor KARRIKIN INSENSITIVE 2 (KAI2), the F-box protein MORE AXILLARY GROWTH2 (MAX2) and the transcriptional repressor SUPPRESSOR OF MAX2 1 (SMAX1), which is ubiquitylated for proteasomal degradation upon KAI2-ligand-induced binding to the KAI2-MAX2 complex. Rice and *Brachypodium distachyon kai*2 mutants are incapable of forming AM. Here, we show that in *Lotus japonicus, Pisum sativum,* and *Nicotiana benthamiana, KAI2* only quantitatively affects AM development, indicating angiosperms vary in their requirement for KAI2-signalling to support AM. Comparative transcriptomics of *L. japonicus* and *B. distachyon* roots after treatment with fungal signalling molecules revealed some AM-relevant genes respond *KAI2*-independently in *L. japonicus* but not in *B. distachyon*. Consistently we obtained evidence for low-level degradation of SMAX1 in *Ljkai2a,b* observed through a ratiometric reporter for the SMAX1 degron (SMAX1_D2_). Further, we found an unexpected accumulation of SMAX1_D2_ in in response to AM even in wild type. Together, this suggests an unexpected role of SMAX1 accumulation in AM roots and that in AM symbiosis of *L. japonicus*, redundant mechanisms drive SMAX1 degradation and gene activation independently of KAI2.

## Introduction

Arbuscular mycorrhiza (AM) is a widespread symbiotic interaction between most land plants and fungi of the *Glomeromycotina* (Spatafora *et al*., 2016; Rimington *et al*., 2020). During this symbiosis, the fungus provides mineral nutrients, such as phosphate and nitrogen, as well as water to the plant, in exchange for photosynthetically fixed carbon in the form of lipids and sugars (Roth & Paszkowski, 2017; Keymer & Gutjahr, 2018; Wipf *et al*., 2019). The development of AM symbiosis is largely controlled by the plant (Gutjahr & Parniske, 2013). Under nutrient-sparse conditions, plants exude a mix of signalling compounds, including carotenoid-derived strigolactones, to recruit AM fungi to their roots (Akiyama *et al*., 2005; Besserer *et al*., 2006). In turn, the fungi release a mix of compounds (germinating spore exudates, GSEs) among which are short-chain chitooligosaccharides, as well as sulphated and non-sulphated lipochitooligosaccharides, together termed ‘Myc-factors’, which are perceived by plant LysM receptor-like kinases (Maillet *et al*., 2011; Buendia *et al*., 2018; Rush *et al*., 2020). At contact with the root surface, the fungus forms a hyphopodium to penetrate the root and makes its way into the cortex. There it colonizes the inside of cortex cells with tree-shaped structures called arbuscules, at which nutrient exchange with the host cell occurs (Bonfante & Genre, 2010). Through a set of signalling proteins together called the Common Symbiosis Signalling Network (CSSN), the plant allows the fungus to enter and grow through the root (Parniske, 2008). In legumes such as *Lotus japonicus* and *Medicago truncatula*, signal transduction following perception of Myc factors (Miyata *et al*., 2014; Carotenuto *et al*., 2017; Zhang *et al*., 2024), as well as rhizobial Nod-factors by LysM receptor-like kinases (Kelly *et al*., 2017), is co-mediated by the malectin-domain containing leucine-rich-repeat receptor-like kinase SYMBIOSIS RECEPTOR-LIKE KINASE (*Lj*SYMRK) (Stracke *et al*., 2002; Antolín-Llovera *et al*., 2014). Unknown events downstream of SYMRK activate perinuclear calcium oscillations, which require the two cation channels *Lj*CASTOR and *Lj*POLLUX, and three cyclic-nucleotide-gated Ca^2+^-channels (CNGC15s) (Charpentier *et al*., 2009, 2016). These calcium oscillations are thought to activate a calcium-and calmodulin-dependent kinase (*Lj*CCamK), which phosphorylates the transcription factor *Lj*CYCLOPS, that interacts with *Lj*DELLA1 (Singh *et al*., 2014; Pimprikar *et al*., 2016). This complex regulates expression of genes necessary for AM symbiosis, such as the GRAS-type transcription factor *Lj*RAM1 (Gobbato *et al*., 2012; Pimprikar *et al*., 2016).

Based on transcriptome profiling, a subset of these genes seems to be controlled by the transcriptional repressor SUPPRESSOR OF MAX1 2 (SMAX1) (Choi *et al*., 2020; Das *et al*., 2025). SMAX1 is the proteolytic target of a hormone-like signalling module referred to as karrikin signalling (Waters & Nelson, 2023). The word ‘karrikin’ refers to a small butenolide molecule, which is generated in the smoke of burnt vegetation, and induces seed germination of fire-following plant species (Flematti *et al*., 2004; Nelson *et al*., 2009). Plants perceive karrikin through the nuclear-cytoplasmically localized alpha/beta-hydrolase protein KARRIKIN INSENSITIVE2 (KAI2) (Sun & Ni, 2011; Waters *et al*., 2012). Following perception, KAI2 interacts with the F-box protein MORE AXILLARY GROWTH2 (MAX2), and similar to other hormone signalling pathways, with the E3-ubiquitin ligase complex ASK1/SKIP1-CULLIN1-F-box (SCF) that polyubiquitinates SMAX1, resulting in proteasomal degradation (Nelson *et al*., 2011; Soundappan *et al*., 2015; Blázquez *et al*., 2020). KAI2 is also proposed to perceive an endogenous ligand, as mutants exhibit a variety of developmental phenotypes independent of post-fire germination (reviewed in Waters & Nelson, 2023; Varshney & Gutjahr, 2023). However, the identity of this compound is still unknown. KAI2 is essential for AM symbiosis in rice, as the fungus does not attach to the root surface of rice *kai2* mutants and consequently does not colonize the root (Gutjahr *et al*., 2015; Choi *et al*., 2020). However, while *kai2* mutants of *Brachypodium distachyon* have a similar phenotype to rice, albeit allowing very low colonization, *kai2* mutants of barley and *Medicago truncatula* are colonized less than wild type but still substantially including the formation of arbuscules (Gutjahr *et al*., 2015; Liu *et al*., 2019; Meng *et al*., 2022; Li *et al*., 2022b). In the liverwort *Marchantia paleacea KAI2* is not required for thallus colonization (Kodama *et al*., 2022). Together this suggests differing requirement of *KAI2* for AM among plant species. Here we interrogated the role of karrikin signalling in AM of the model legume *Lotus japonicus* as well as in pea and *Nicotiana benthamiana* and found that KAI2 has a quantitatively smaller impact in these species as compared to rice and *B. distachyon*.

## Methods

### Plant material, growth conditions and AM assays

*Lotus japonicus* ecotype Gifu was used as wild type and background for mutations. Isolation of *L. japonicus d14-1, max2-1, max2-3, max2-4, kai2a-1, kai2b-1, kai2b-2*, *kai2b-3* mutant lines through LORE1 transposon insertion or TILLING was described previously (Carbonnel *et al*., 2020b). For germination, *L. japonicus* seeds were scarified with sand-paper and surface sterilized with 1% NaClO and 0.1% SDS and then washed with MilliQ water for 5 times. Imbibed seeds were germinated on 0.8% Plant Agar (Duchefa, Netherlands) at 24 °C in light. Arbuscular Mycorrhiza experiments were performed either in open pots, cones or in PTC containers. Experiments in PTC containers (Duchefa, Netherlands) were performed as previously described in (Torabi *et al*., 2021). In short, 4 seedlings (5-7 days old) were planted with 400-500 spores per plant *R. irregularis* DAOM 197198 (type C, Agronutrion, Toulouse, France) in PTC containers filled with 300 g quartz sand and watered once with 30 mL of ½ strength Hoagland containing 25 µM phosphate. After planting, the plants were grown under a long-day photoperiod (16 h light at 23 °C/8 h dark at 21 °C) at a light intensity of 109 µE. In open pot experiments, *L. japonicus* seedlings (7 – 14 days) were planted into quartz sand with 400 spores of *R. irregularis* and watered twice a week with 25 mL ½ strength Hoagland for the first two weeks, followed by biweekly watering with nutrient solution and an additional weekly watering with deionized water.

For *Pisum sativum* colonization assays, wild type *Cameor*, *kai2a*, *kai2b*, and *kai2a kai2b* mutant seeds were obtained from Alexandre de Saint Germain (INRA Versailles, Guercio *et al*., 2022). Seeds were surface sterilized with 1% NaClO and 0.1% SDS and washed with milliQ water for 5 times. Seeds were pre-germinated on 0.8% Plant Agar (Duchefa, Netherlands) at 24 °C in light. Then they were planted into open pots with a 2:1 Quarz sand:vermiculite mix with 1000 *R. irregularis* spores per plant. Plants were watered twice a week with ½ strength Hoagland for the first two weeks, followed by biweekly watering with nutrient solution and an additional weekly watering with deionized water.

For the *Nicotiana benthamiana* experiments, seeds were surface sterilized with 1% NaClO and 0.1% SDS, washed with MilliQ water 5 times and germinated on 0.8% plant agar. 7-14 day-old seedlings were planted in quartz sand/vermiculite (1:2) mixture with 500 spores *R. irregularis* per plant, and watered biweekly with B&D medium (Torabi *et al*., 2021) (50 µM phosphate, 7.5 mM nitrogen) and once a week with deionized water. After harvesting, roots were cleared in 10% KOH at 95 °C for 10 minutes (*L. japonicus*) or 25 minutes (*N. benthamiana*), washed in 10% Acetic acid and then stained in 5% Black ink (Pelikan 4001, *Brilliant Black*), 5% Acetic acid. Root length colonization (RLC) of the roots was assessed using bright field microscopy according to an adjusted grid-section intersect method (Torabi et al 2021).

### Construct generation and hairy root transformation

gDNA and promoter sequences of *KAI2b, MAX2* and *SMAX1* were amplified using Phusion^®^ High-Fidelity DNA polymerase (NEB, USA, Cat No. F530S) and ligated into a bacterial Golden Gate LI vector and subsequently moved into multi-gene LIII constructs, including an mCherry transformation marker, as described before (Table S1 (Binder *et al*., 2014)). *L. japonicus* hairy roots were transformed with these LIII constructs using *Agrobacterium rhizogenes*, as previously described (Takeda *et al*., 2009). Transformed roots were screened under a stereo microscope (Leica MZ16 FA; Leica, Germany) to select roots with mCherry fluorescence. For comparison of transformed and non-transformed roots, roots from the same root system but without mCherry fluorescence were also collected.

### Cas9-mediated generation of quadruple *Nbkai2a,b,c,d* mutant

Four KAI2 paralogs were found in the *N. benthamiana* genome by blasting the protein sequence of *At*KAI2 against the predicted proteome of *N. benthamiana* (SolGenomics): Niben101Scf23299g00008 (*KAI2a*), Niben101Scf03009g01006 (*KAI2b*), Niben101Scf04262g00025 (*KAI2c),* and Niben101Scf11178g00001 *(KAI2d)*. Of these, *KAI2c* is a pseudogene with an early stop codon in exon 2 (Fig. S1). 4 gRNAs (Table S2) were designed using DESKGEN online platform (Doench *et al*., 2016) to target the three genes and cloned using the CasCADE modular vector system (Hoffie, 2022; Rezaeva *et al*., 2025) to generate pGH634. Briefly, forward and reverse DNA oligonucleotides (Table S3) were generated for gRNA containing overhangs for BbsI-based insertion into the gRNA module vectors pIK75 to pIK78 containing the Arabidopsis *U6* promoter (U6-26-p). The four individual gRNA module vectors (pGH743-pGH746) were assembled using the backbone vector pIK19 to generate the gRNA assembly vector pGH747. This gRNA assembly was then combined with the *cas9* endonuclease expression unit (pBR20) and an auxiliary unit (pIK155). The intermediate assembly vector pGH747 contained four gRNA expression units and an Arabidopsis codon-optimised *cas9* under the parsley *UBIQUITIN 4-2* promoter (*PcUBI4-2*). Finally, the genome editing modules of pGH747 were cloned into the generic binary vector p6i-d35S-TE9 (DNA-Cloning-Service e.K., Hamburg, Germany) via SfiI restriction and ligation. The binary vector pGH634 also contains the *hpt* gene under the control of a doubled-enhanced *CaMV35S* promoter for plant selection. The final plasmid was verified through Sanger sequencing and introduced into *Agrobacterium tumefaciens* GV2260 via electroporation. Due to sequence duplication across *KAI2c and KAI2d,* gRNA 1 also targeted the presumed non-functional pseudogene *KAI2c*. Wild-type *N. benthamiana* seedlings underwent transformation with pGH643, and T1 plants were screened for mutations in the regions containing the four *KAI2* homologs. T2 plants were screened for the absence of the transgene, and T3 plants was screened for deletions causing a non-functionality of genes, by PCR amplification with primers (Table S3) spanning the regions targeted by the respective gRNAs, and subsequent incorporation into L1 Golden Gate plasmids and Sanger sequencing (Binder *et al*., 2014).

### Confocal microscopy and quantitative analysis in Fiji

For arbuscule imaging, roots were harvested and kept in 80% ethanol at 4°C prior to imaging. For staining with wheat germ agglutinin (WGA)-AlexaFluor488, plants were cleared in 20% KOH for 2 days at room temperature, then incubated with 0.3% HCl for 2 hours and subsequently washed with ddH_2_O and then with 1x PBS buffer (pH 7.4) three times each. Roots were then incubated with WGA-AlexaFluor488 at 0.4 µg/mL and imaged using a Leica Stellaris 8 confocal microscope (exc. 491 nm; emission 519 nm). If used for sectioning, roots were harvested, checked for transgene activity by fluorescence under a fluorescence stereomicroscope (MZ10F, Leica), and pieces of multiple roots were mounted in parallel in 4% agarose in 1x PBS (pH 7.4) for sectioning with a vibratome. Sections were stained with SCRI Renaissance Stain (RS2200; Renaissance Chemicals) for at least 10 minutes and then imaged using a Leica Stellaris 8 confocal microscope. For imaging SMAX1_D2_-mScarlet-I/Venus, laser intensity and gain were kept equal for all images, and images were obtained by sequential scanning (Venus: exc. 523 nm, emm. 571 nm; gain, 2.5; laser intensity at 5.59%. mScarlet-I: exc. 576 nm emm. 638 nm; gain 58.3, laser intensity at 9.20%. SR2200: exc. 420 nm, emm. 482 nm; gain 2.5; laser intensity at 2.5%). For quantitative fluorescence analysis, background was removed in Fiji with a rolling ball radius of 50 pixels using sliding paraboloid function. Nucleus circumferences were traced manually, and the mean fluorescence intensity of the so-defined ROIs was measured for the Venus and mScarlet-I channel. For statistical comparison of mScarlet-I/Venus, ratio across conditions, genotypes and tissue types, only nuclei with a minimum intensity of 350 a.u. for the Venus channel were included. Prior to this decision, different cut-offs were consulted, and the order of median values across respective remained the same regardless of the cut-off (Table S4). For the statistical analysis, a linear mixed-effects model on log-transformed data, including log(Venus) as a covariate, was used.

### Production of *R. irregularis* germinating spore exudates

Spores of AM fungi (type A, Agronutrion, Toulouse, France) were washed 3 times in ½ Hoagland and consequently incubated in minimal M medium (Hildebrandt *et al*., 2002) for 1 week at 28°C. The solution was centrifuged, and the supernatant containing GSEs but no spores or fungal hyphae was removed and stored at-70°C. For treatment, the mixture was diluted in ½ Hoagland medium to a level of GSE from 210 spores/mL for plant treatment.

### RNA sequencing and transcriptome analysis

#### RNA sequencing

For RNA-seq after treatment with chitotetraose (CO4) or germinating spore exudates (GSEs), 5-day-old *L. japonicus-* and *B. distachyon-*seedlings were grown in PTC (Torabi *et al*., 2021), but without *R. irregularis* spores. After 3 weeks, pots were opened for 24 hours and treated with 10 nM CO4 (Megazyme, Ireland) or GSEs harvested from *R. irregularis* spores for 6 hours. Roots were then harvested in liquid nitrogen and stored at-70°C. RNA was extracted using the Spectrum™ Plant Total RNA Kit (Sigma-Aldrich, USA). RNA samples were treated with Turbo DNase (Thermo Scientific, USA, Cat No. AM1907). Sample concentration was quantified using a NanoDrop (Thermo Scientific, USA), and RNA quality was assessed using a Bioanalyzer (Agilent, USA). Samples with RIN values of 7 or higher were used for library preparation using a QuantSeq 3’ library kit (Lexogen, USA) and sequenced on Illumina HiSeq 2500 to obtain 100 bp single-end reads (as previously described in Das *et al*., 2025).

#### Quality control and read trimming

Raw sequencing reads from 72 RNA-seq libraries (36 *B. distachyon*, 36 *L. japonicus*) were assessed for quality using FastQC (v0.12.1; Andrews, 2010). Adapter sequences and low-quality bases were removed using Trimmomatic (v0.39; Bolger *et al*., 2014) in single-end mode with the following parameters: ILLUMINACLIP:2:30:10, LEADING:3, TRAILING:3, SLIDINGWINDOW:4:15, MINLEN:40, using a custom adapter file. Trimmed reads were re-evaluated with FastQC.

#### Read alignment

Trimmed reads were aligned to their respective reference genomes using HISAT2 (v2.0.6; Kim *et al*., 2019) (v2.0.6). For *B. distachyon*, reads were mapped to the v3.0 assembly (annotation v3.2, Phytozome; The International Brachypodium Initiative, 2010). For *L. japonicus*, reads were mapped to the Gifu v1.2 genome assembly (LotusBase; Kamal *et al*., 2020). Alignments were performed using default HISAT2 parameters with per-sample read-group metadata. Resulting alignments were coordinate-sorted and indexed in csi format using SAMtools (v1.1.3; Li *et al*., 2009).

#### Read quantification

Gene-level read counts were generated using *featureCounts* (subread v2.0.1; Liao *et al*., 2014) with modified GFF3 annotation files. To improve read assignment for 3′ mRNA sequencing, a custom script was used to extend the last exon of each gene by up to 3 kb or until the next gene. For *B. distachyon*, a modified version of the *Bdistachyon_556_v3.2* gene annotation was used, while for *L. japonicus*, a modified Gifu v1.3 annotation was applied, excluding chloroplast and mitochondrial genes. Reads were assigned to genes based on the ID attribute at the gene feature level, with strand specificity enforced (–s 1). Multi-mapping reads were retained and reads overlapping multiple features were counted for all relevant features (–M –O), minimizing read loss associated with the 3′ mRNA sequencing and exon extension strategy.

#### Differential expression analysis

Prior to differential expression analysis, five *B. distachyon* samples (Bdi_41_WT_CO4_S8, Bdi_7_kai2_Mock_S19, Bdi_8_kai2_Mock_S20, Bdi_9_kai2_Mock_S21, Bdi_31_kai2_CO4_S25; Fig. S2) were identified as outliers based on Euclidean distance from group centroids in PCA space exceeding the 95th percentile of the within-group distribution and hierarchical clustering of sample-to-sample distances. These samples were excluded prior to further analysis. No samples needed to be excluded for *L. japonicus* (Fig. S3). Differential expression analysis was performed independently for each species using DESeq2 (v1.36.0); (Love *et al*., 2014) with a model design of ∼ 0 + Group, where Group encoded the combination of genotype (*kai2* mutant or wild type) and treatment (mock, CO4, or GSE). Genes with fewer than 10 summed counts across all samples were excluded prior to model fitting. Nine pre-specified pairwise contrasts were tested, capturing the genotype effect without treatment, treatment effects within each genetic background, and genotype-by-treatment interactions. Multiple testing correction was performed using the Benjamini-Hochberg procedure. Genes were considered differentially expressed at an adjusted p-value ≤ 0.05 an no fold-change cut-off.

#### Expression pattern visualization and clustering

Variance-stabilizing transformation (VST, blind = FALSE) was applied to filtered count matrices. Per-gene z-scores were computed across all samples and the median z-score per treatment group was used as a summary statistic. The union of DEGs from five key contrasts was used for clustering. Hierarchical clustering was performed using complete linkage on Euclidean distances. The optimal number of clusters was determined after inspecting elbow and average silhouette score analysis across k = 2–25 (random seed 369), yielding k = 7 for *L. japonicus* and k = 5 for *B. distachyon*. Expression heatmaps were generated using pheatmap (Kolde, 2025).

#### Orthology analysis

Orthologous gene groups were inferred using OrthoFinder (v2.5.5; (Emms & Kelly, 2019)) run with multiple sequence alignment mode using MUSCLE (-M msa-A muscle-t 32; Edgar, 2004) on the primary predicted proteomes of *B. distachyon* (Bdistachyon_556_v3.2, primary transcripts only) and *L. japonicus* (Gifu v1.3, longest isoforms). DEGs from each contrast and species were mapped to their orthogroups and overlap between species was visualized as UpSet plots using ComplexUpset (R) (https://krassowski.github.io/complex-upset/) and upsetplot (Python) (Lex *et al*., 2014).

### RNA extraction and RT-qPCR

RNA was extracted using the RNA isolation using Spectrum™ Plant Total RNA Kit (Sigma-Aldrich, USA). DNase treatment and subsequent cDNA synthesis was performed using the *BIO-RAD* iScript^TM^ gDNA clear cDNA Synthesis kit using 600 ng of RNA for every sample. cDNA was diluted ten times, and qPCR was run using diluted cDNA in the Jena Bioscience SYBR green kit in a volume of 7 µL in 384-well plates (Roche) using the Roche LightCycler 480 Instrument II.

## Results

### KAI2 has a quantitative effect on root length colonization across several dicotyledon species

KAI2 plays an important role in AM symbiosis in rice and *B. distachyon*, as *d14l/kai2* loss-of-function mutants abolish or exhibit a drastic decrease in root colonization by AM fungi (Gutjahr *et al*., 2015; Choi *et al*., 2020; Meng *et al*., 2022; Hong *et al*., 2026). To investigate whether this role for karrikin signalling in AM symbiosis is conserved across angiosperms, we characterized root colonization by *Rhizophagus irregularis* of *L. japonicus* wild type, *kai2a, kai2b, kai2a kai2b* (from here on *kai2a,b*), *d14,* and *max2* plants (Fig. 1a; Fig. S4a-b). Compared to wild type, *max2* and *kai2a,*b double mutants exhibited a significant approximately 50% decrease in root length colonization (RLC) at 42 days post inoculation (dpi), whereas RLC was not changed in *d14*, *kai2a* or *kai2b* single mutants (Fig. 1a; Fig. S4a). This indicates that in the context of AM symbiosis, *KAI2a* and *KAI2b* function redundantly, and that strigolactone signalling does not regulate RLC in *L. japonicus*. The morphology of all intraradical fungal structures, including arbuscules in *kai2a,b* or *max2* mutant roots appeared like in the wild type (Fig. S5) and the RLC by intraradical hyphae, arbuscules and vesicles recapitulated total RLC (Fig. S4b). Therefore, for the rest of this study, we focused on total RLC. Transgenic complementation of *kai2a,b* and *max2* mutant hairy roots with either a *KAI2b_pro_:gKAI2b-myc*, or *MAX2_pro_:gMAX2* expression cassette, respectively, resulted in recovery of wild-type levels of RLC for roots containing the respective transgene (Fig. 1b-c), while non-transformed roots of the same root systems maintained lower colonization levels, even when all genotypes were grown in the same pot (Fig. 1b). Together, this indicates that in *L. japonicus*, disruption of karrikin signalling results in a quantitative decrease in RLC and that *KAI2* and *MAX2* are locally and likely not systemically required to promote AM. The AM phenotype of *L. japonicus kai2a,b* and *max2* is much less severe than previously found for rice and *B. distachyon kai2* mutants (Gutjahr *et al*., 2015; Meng *et al*., 2022; Hong *et al*., 2026), indicating a reduced requirement of the KAI2-MAX2 signalling complex for AM symbiosis in *L. japonicus*.

**Fig. 1.**
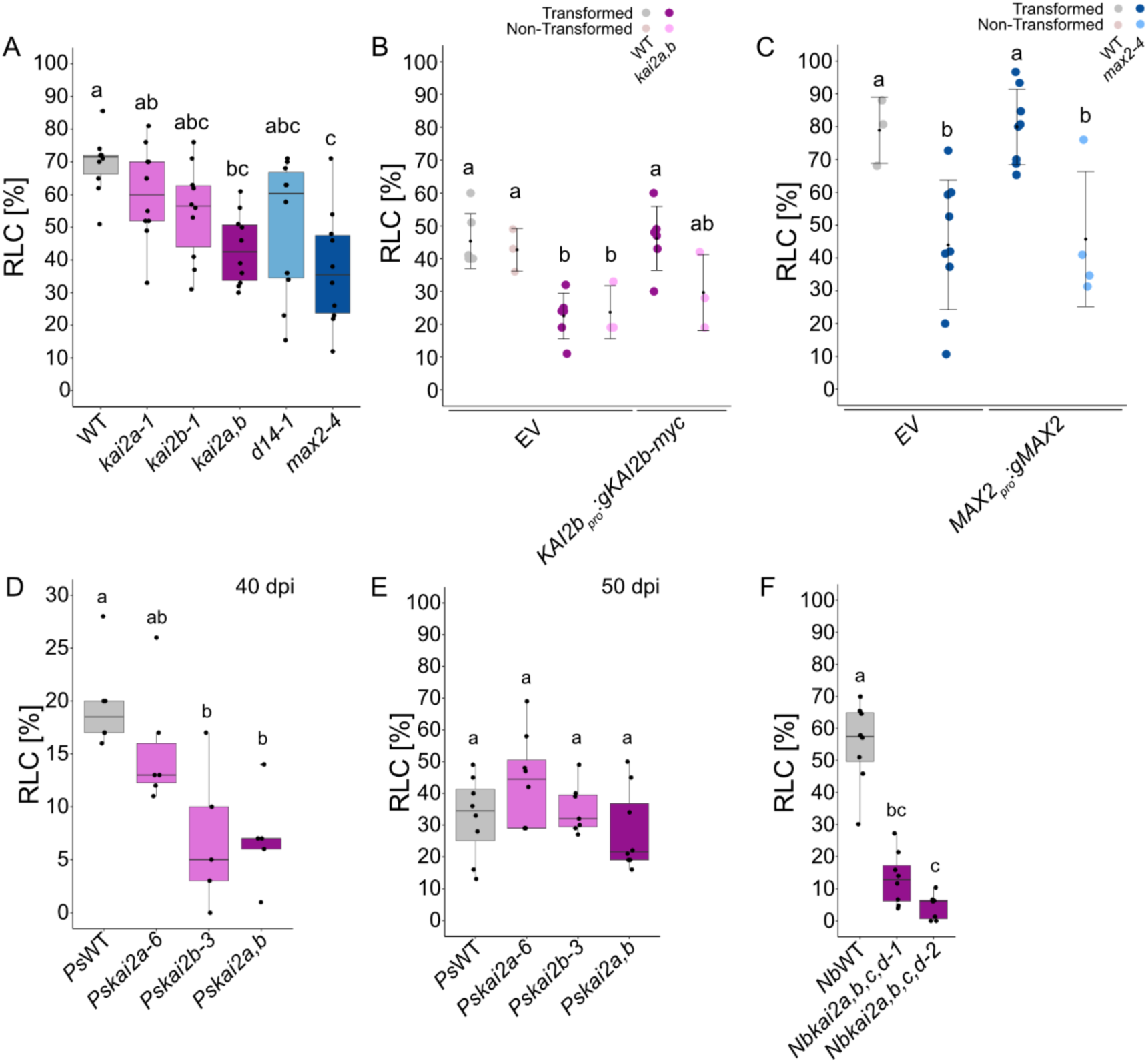
Karrikin receptor mutants in legumes show a quantitative reduction of root length colonization. Percent total root length colonization (RLC) by *Rhizophagus irregularis* of (a) *L. japonicus* strigolactone and karrikin perception mutants grown in sand-vermiculite open pot cultures for 42 dpi. N_plants_ = 8-10 ANOVA, Tukey, p<0.05, F_5/54_=5.862; (b) *L. japonicus* wild type and *kai2a,b* mutants growing transgenic (transformed) and non-transgenic (non-transformed) roots after *Agrobacterium rhizogenes*-mediated transformation with empty vector (EV) or *pKAI2b:KAI2b-Myc* grown in competition in the same sand and closed plant tissue culture containers (PTC) for 21 days post inoculation. ANOVA, Tukey, p<0.05, F_5/22_= 8.528. (c) *L. japonicus* wild type and *max2-4* plants producing transgenic and non-transgenic roots after transformation with an EV or *pMAX2:MAX2* expression cassette and grown in sand vermiculite open pot cultures for 42 dpi. N_plants_ = 6-8, ANOVA, Tukey, p<0.05, F_3/20_= 8.87; (d-e) *Pisum sativum* wild type and *kai2* single and double mutants grown in sand vermiculite open pot cultures for (d) 40 and (e) 50 dpi. N_plants_ = 6-8, ANOVA, Tukey, p <0.05, F_3/84_=8.17; (f) *Nicotiana benthamiana* wild-type and *kai2a,b,c,d* quadruple mutants grown in sand vermiculite open pot cultures for 35 dpi. N_plants_ = 7-8 Kruskal-Wallis, Dunn test, p < 0.05. For figures (a), (d), (e), (f), bold horizontal lines represent median value; lower and upper whisker indicate quartile 1 and quartile 4. Lower and upper horizontal lines indicate minima and maxima. For figures (b) and (c), black dot indicates mean value, whiskers error bars indicate standard deviation of the mean. In all graphs, different letters indicate different statistical groups. All experiments were repeated at least 2 times with similar results.

To examine whether this is the same for other legume species, we tested root colonization by *R. irregularis* of *Pisum sativum* wild type, *kai2a, kai2b* and *kai2a kai2b* (from here on *Pskai2a,b*; Fig. 1d-e). At 40 dpi, *Pskai2a,b*, similar to *Ljkai2a,b*, showed a relatively mild (approx. 15%) reduction in RLC, compared to wild type. However, at 50 dpi RLC of *Pskai2a,b* caught up with the wild type, and the quantitative difference was no longer visible (Fig. 1d-e). These findings, together with results from *M. truncatula* (Li *et al*., 2022b), indicate the requirement of *KAI2* for AM symbiosis is reduced in legumes as compared to rice and *B. distachyon*, and that in *P. sativum, KAI2* plays its main role during the earlier phase of root colonization (Fig. 1e). In addition, to their AM-competence, legumes evolved the ability to establish root-nodule symbiosis (RNS) with rhizobia. In order to host these nitrogen-fixing bacteria, legumes have co-opted genes from the CSSN, which are also of fundamental importance for AM (Parniske, 2008). Karrikin signalling has been shown to influence expression of some genes that are part of the CSSN in rice and in *L. japonicus* roots (Choi *et al*., 2020; Das *et al*., 2025; Hong *et al*., 2026), but it does not appear to play a role in RNS in *M. truncatula* (Li *et al*., 2022b). Therefore, CSSN-genes may be transcriptionally regulated by multiple independent pathways in legumes, which may explain the reduced requirement of KAI2 for AM symbiosis in this clade. To understand whether this reduced requirement of *KAI2* for AM symbiosis is specific to legumes or also observed for dicotyledons that do not form RNS, we generated a quadruple *kai2a,b,c,d* knock-out mutants in *Nicotiana benthamiana* by Cas9 editing (Fig. S1). Two allelic quadruple *Nbkai2a,b,c,d* knock-out lines exhibited a reduction in RLC compared to wild-type roots (Fig. 1f). Although the reduction in RLC in *Nbkai2a,b,c,d* roots seems stronger than in *Ljkai2a,b* or *Pskai2a,b*, some *Nbka2a,b,c,d* plants still reach up to 25% RLC, thus much higher than previously described for *d14l* in rice (0% RLC) and *kai2* in *B. distachyon* (up to 10% RLC; Gutjahr *et al*., 2015; Choi *et al*., 2020; Meng *et al*., 2022). This indicates that the indispensability of *KAI2* for AM symbiosis is not conserved across vascular plants and may be specific to certain species. Indeed, a previous study even reported for another monocotyledon barley that *d14l*/*kai2* roots were colonized by approximately 10-20% RLC (approximately 60-70% reduction with respect to wild type) (Li *et al*., 2022b), indicating that not only in dicotyledons but also in some poaceous monocotyledons *KAI2* impacts AM mainly quantitatively.

### Loss of *KAI2* divergently affects transcriptional responses to CO4 and GSEs in *L. japonicus* vs. *B. distachyon* roots

To assess how loss of *KAI2* affects the response of roots to AM fungi, we used the transcriptome as a marker and performed RNA-sequencing on roots of *L. japonicus* and *B. distachyon* wild-type and *Bdkai2/Ljkai2a,b* roots. Colonized *kai2* roots are not comparable between *L. japonicus* and *B. distachyon*, as *kai2* mutants of both species show drastically different RLC (Meng *et al*., 2022, this work). This biases any transcriptome difference towards genes that respond cell-autonomously to colonization causing their expression to be correlated with RLC. Therefore, we treated roots with signals released by fungal spores during germination, namely chitotetraose (CO4) and the complex mixture of germinating spore exudates (GSEs). An initial exploratory analysis using Padj < 0.05 and no Fold-Change cut-off (because the gene induction amplitude to AM fungal signals is usually low) identified 2537 unique differentially expressed genes (DEGs) for *L. japonicus*, and 809 DEGs for *B. distachyon* (Fig. S6a-b, Table S4). In *L. japonicus,* the majority of DEGs (91%, 2303 DEGs) were found when *kai2a,b* was compared to wild type in the absence of treatment (Fig. S6a, cluster 2 & 3; Table S5). In *B. distachyon,* this comparison comprises a smaller proportion of the total amount of DEGs (29%, 235 DEGs; Fig. S6b, cluster 6; Table S5). This indicates that for *L. japonicus*, loss of *KAI2* seems to have a stronger effect on the transcriptome of non-treated roots than for *B. distachyon*. The relatively low amount of DEGs between *Bdkai2* and the corresponding wild type is consistent with a previous study that identified 123 DEGs for the comparison of entire wild-type and *Bdkai2* seedlings (Meng *et al*., 2022).

Also, the number of DEGs in response to CO4 and GSE differed between *L. japonicus* and *B. distachyon* (Fig. 2a-b). Overall, wild type *L. japonicus* roots responded with more genes to CO4 than to GSE treatment (115 vs 68 DEGs), whereas wild-type *B. distachyon* roots were virtually unresponsive to CO4 treatment, but showed a strong response to GSE-treatment (14 vs 362 DEGs; Fig. 2a-b). This divergence is likely caused by different LysM receptor repertoires of the two species. Interestingly, *kai2* mutants of both species barely responded to GSE-treatment (Fig. 2a-b), indicating that the transcriptional response to GSEs (at least at 6h post-treatment) requires *KAI2*. However, CO4 treatment of *L. japonicus* roots resulted in a similar amount of DEGs for wild type and *kai2a,b* (115 DEGs for wild-type, 101 for *kai2a,b;* Fig 2a). Moreover, the Log2FC of these DEGs for CO4 vs mock was very similar between wild type and *kai2a,b* (Fig. 2c, clusters 1 and 2). Conversely, in *B. distachyon,* wild-type and *kai2* roots display a contrasting transcriptional response to CO4 treatment with wild type showing little to no response (16 DEGs in CO4 over mock), and *kai2* differentially regulating almost 20-times more genes in response to CO4 treatment (208 DEGs for CO4 vs. mock; Fig. 2b). Interestingly, more than half of these 208 genes are downregulated after CO4 treatment (137 genes, cluster 3 of Fig. 2d), which is in sharp contrast with the majority of DEGs being upregulated in response to CO4 in *Ljkai2a,b* (Fig. 2c-d). To understand whether there is functional overlap in the transcriptional response of the two species, we ran an Orthofinder analysis to identify orthologs across *L. japonicus* and *B. distachyon* (Fig. S6). However, there were little to no overlaps in orthogroups across treatments and genotypes for either CO4 or GSE treatment (Fig. S6C, D). This indicates clear differences in the response to Myc-factor perception between *L. japonicus* and *B. distachyon* for both CO4 and GSE treatments, and in how loss of *KAI2* function affects the transcriptional response to CO4 treatment. To understand whether genes in *L. japonicus* that are upregulated in response to CO4 in wild type and *Ljkai2a,b* are affected by karrikin signalling, we compared them to genes with increased expression in untreated *smax1* mutant vs. wild-type roots from a previous study (Fig. 2e, (Das *et al*., 2025)). CO4-treated *kai2a,b* roots exhibit 37 DEGs overlapping with *smax1* of which 19 also overlap with CO4-induced genes in wild type (Fig. 2e), indicating that despite *KAI2* loss-of-function, genes directly or indirectly suppressed by SMAX1 can still be induced by CO4, potentially through SMAX1-independent transcriptional regulators or due to redundant pathways affecting SMAX1 accumulation (e. g. the strigolactone receptor D14, (Li *et al*., 2022a)). Among the genes with increased expression after CO4 treatment and in *smax1* mutants were *CCaMK* (in wild-type)*, MAX1* and *VAPYRIN1* (in *kai2a,b*), and *DXS2* and *NOPE1* (in both wild type and *kai2a,b*) known to be key regulators of symbiosis, involved in strigolactone biosynthesis, intracellular accommodation of the fungus, the biosynthesis of a mevalonate precursor, and rhizosphere communication from plant to fungus through N-acetylglucosamine derivatives (Walter *et al*., 2002; Floß *et al*., 2008; Pumplin *et al*., 2010; Cardoso *et al*., 2014; Nadal *et al*., 2017; Liu *et al*., 2021). Also, *ERN1* was induced both by CO4 treatment in wild type and *kai2a,b* and in *smax1* mutant roots (Fig. 2e). *ERN1* encodes an AP2-domain Ethylene Response Factor with a known role in nodulation, but has not been described to be important for AM symbiosis, although it was found to be upregulated in AM colonized tomato-roots (Cerri *et al*., 2017; Stuer *et al*., 2026). The overlap with genes showing increased expression in *smax1* seems to be specific for CO4 treatment, as genes induced by GSE-treatment in *L. japonicus* did not show such a substantial overlap (Fig. S7c).

**Fig. 2.**
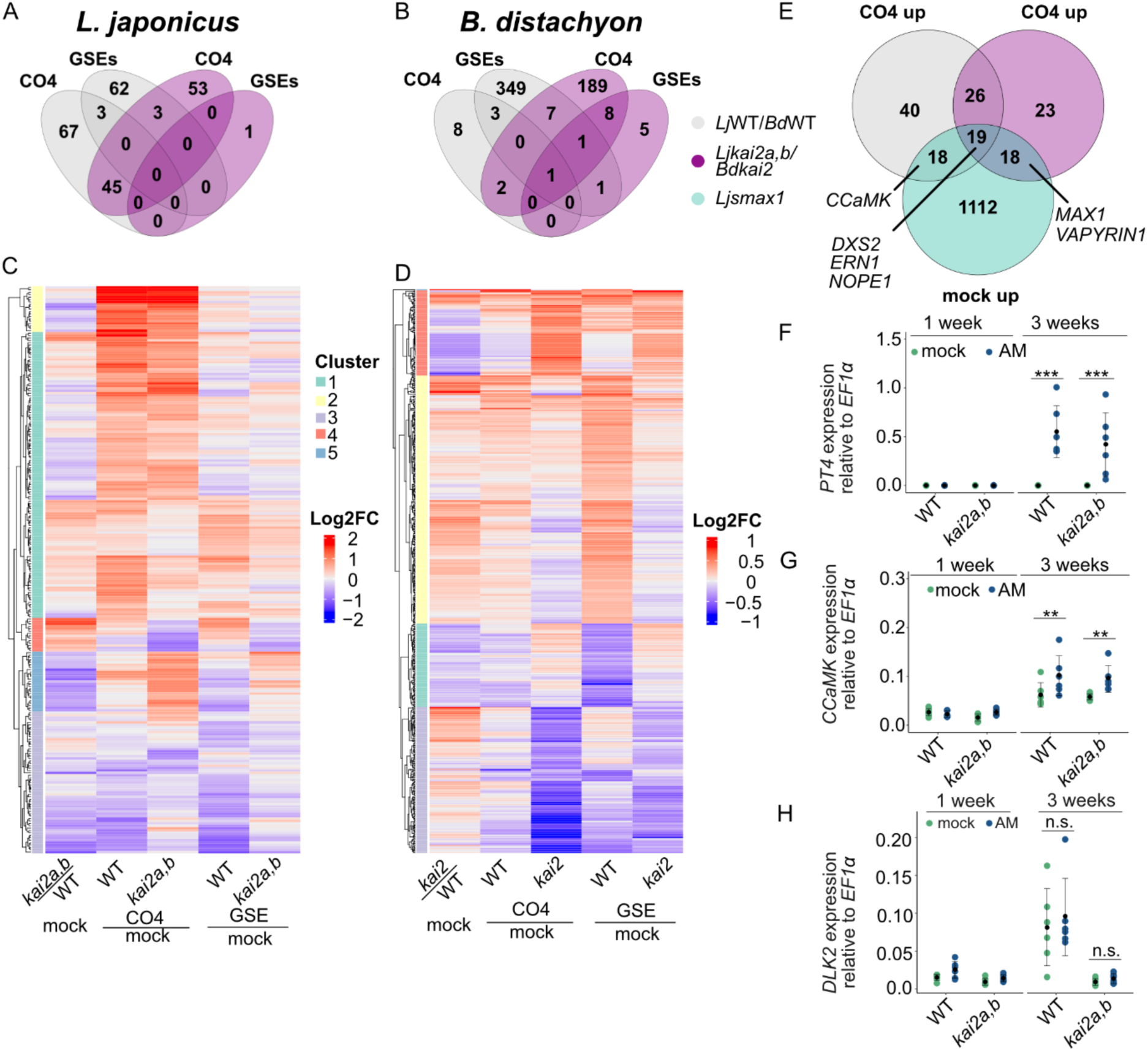
*L. japonicus* and *B. distachyon* exhibit distinct transcriptional responses after treatment with fungal signals. (a-b) Venn diagrams showing numbers of differentially expressed genes (padj < 0.05) in roots after a 6-hour treatment with CO4 or GSE across the indicated genotypes in (a) *L. japonicus* or (b) *B. distachyon* grown in closed PTC-containers for 3 weeks before treatment. (c-d) K-means clustered (k = 5 for both species) heatmaps visualizing Log_2_FC of genes in (a) or (b) for CO4 or GSE over mock treatment in wild type (WT) or *kai2* mutant roots, or in *kai2* mutant over WT under mock conditions for (c) *L. japonicus* or (d) *B. distachyon* roots. (e) Venn diagram comparing genes differentially expressed upon CO4 treatment in *L. japonicus* WT or *kai2a,b* (this work), and in *smax1* compared to WT roots in mock conditions (from Das *et al*. 2025). Relative expression (to *LjEF1α*) of (f) the AM symbiosis marker gene *LjPT4,* (g) *LjCCaMK*, and (h) *LjDLK2* in roots of *L. japonicus* WT or *kai2a,b*, at 1 or 3 wpi without (mock) or with spores of *R. irregularis* (AM). Asterisks indicate statistically significant differences. N_samples_= 6, 2 root systems per sample, 3-way ANOVA, estimated marginal means contrast for AM treatment, * - p<0.05; ** - p<0.01, *** - p< 0.001. F-value for treatment:harvest interaction; f) F_1/40_ = 31.997, g) F_1/40_ = 8.709, h) F_1/40_ = 0.008. Black dots represent mean values, bars represent standard deviation of the mean.

We also visualized the Log2FC of genes known to be involved in AM symbiosis, karrikin signalling, and strigolactone biosynthesis and signalling (Fig. S7a-b). This revealed a subset of AM-relevant genes with reduced expression in *L. japonicus kai2a,b* as compared to wild type (such as *CYCLOPS*, *Zaxinone synthase ZAS1* and *2, NSP1, MAX1*). Some were clearly induced by CO4 (but not GSE) treatment in roots of both genotypes (such as *DXS2* and *ERN1*) (Fig. S7a). Overall, in *B. distachyon*, the effect of *KAI2* mutation on the transcriptome is less pronounced, but multiple genes, including *ERN1* and the GRAS transcription factor gene *NSP2* with roles in AM symbiosis, regulation of strigolactone biosynthesis and nodulation (Liu *et al*., 2011; Cerri *et al*., 2017; Li *et al*., 2022b; Yuan *et al*., 2023; Hong *et al*., 2026) are expressed at a lower level in *kai2*-mutants (Fig. S7b). Interestingly, CO4-treatment results in a significant downregulation of *KAI2* expression in *Bdkai2*. Because the *Bdkai2-2* allele carries only a point mutation, the gene is still expressed (Meng *et al*., 2022). Overall, few orthologous genes are similarly deregulated in untreated *kai2* mutants of both *L. japonicus* and *B. distachyon*. The exception is the karrikin signalling marker gene *LjDLK2*/*BdDLK2b,* which is significantly less expressed in both *Ljkai2a,b* and *Bdkai2* compared to the respective wild types (Fig. S7a-b). To examine, if genes responding to CO4 treatment in WT and/or *kai2a,b* roots also respond to root colonization by AM fungi, we determined expression of the common symbiosis gene *Calcium Calmodulin dependent Kinase* (*CCaMK*), as a marker along with the AM-marker gene *PHOSPHATE TRANSPORTER4* (*PT4*) at an early and later time point of colonization using RT-qPCR (Fig. 2f-g). *PT4* was not detected at 1-week post-inoculation (wpi) in either genotype regardless of the presence of fungal spores, indicating that no arbuscules had formed at this time point. At 3 wpi, it was well expressed in both inoculated wild-type and *kai2a,b* roots, indicating colonization (Fig. 2f). *CCaMK* expression was slightly but significantly increased upon root colonization in wild-type and *kai2a,b* roots (Fig. 2g). Additionally, in *kai2a,b* roots, we detected a significant increase in *MAX1* expression at 3 wpi (Fig. S7d). This indicates that the upregulation of certain AM-relevant genes in *Ljkai2a,b* occurs not only after CO4 treatment, but also during root colonization and suggests that in *L. japonicus KAI2* is not required for this induction, for example, due to redundant mechanisms. The low level of induction detected in mRNA from bulk tissue is likely due to dilution because it probably occurs cell-specifically. In contrast, *DWARF14-LIKE2* (*DLK2)*, a gene encoding an alpha/beta-hydrolase homologous to *KAI2* and *D14,* used as a marker gene for activation of karrikin signalling through proteasomal degradation of SMAX1 (Nelson *et al*., 2010, 2011; Waters *et al*., 2012) was not induced in colonized *kai2a,b* roots (Fig. 2h). This suggests regulatory redundancy for some but not all putative SMAX1 downstream targets.

### Karrikin signalling regulates fungal spread within roots of *L. japonicus*

In rice, mutation of KAI2 results in absence of root colonization, while mutation of the transcriptional repressor *SMAX1* results in increased RLC. Similarly, *L. japonicus smax1* mutants exhibit an increase in RLC with *R. irregularis* compared to wild type (Fig. 3a; Das *et al*., 2025). *Ljkai2a,b smax1* triple mutants exhibit a similar increase, indicating the function of *SMAX1* is epistatic to *KAI2a* and *KAI2b* (Fig. 3a), as previously described for rice (Choi *et al*., 2020). This increased RLC persisted over time. While colonization also increased over time in *kai2a,b* mutants, their RLC remained lower than in the wild type (Fig. 3b). Expression of *SMAX1_pro_:HA-SMAX1* in *smax1* hairy roots restored RLC to wild-type levels (Fig. S8). To inspect whether the increased colonization of *smax1* roots is caused by an increase in infection points or increased fungal spread within the roots, we quantified the number of colonization units (CUs; fungal intraradical unit, originating from one root entry) and the length of CUs per plant in wild-type, *kai2a,b* and *smax1* roots. Roots of *kai2a,b* contained less CUs than both wild-type and *smax1* roots, which displayed similar CU numbers (Fig. 3c). However, CUs in *smax1* roots were on average longer than CUs in *kai2a,b* roots, while CUs in wild-type roots had an intermediate length (Fig. 3d). This indicates that the increased levels of RLC in *smax1* roots is most likely due to a faster fungal spread through the roots, rather than an increase in infection points. Karrikin signalling also seems to affect the morphology of CUs, with shorter compact CUs in *kai2a,b* roots, and long hyphae with seemingly lower arbuscule density in roots of *smax1* at early stages of colonization (Fig. 3E). However, arbuscules in *smax1* roots were fully branched and similar to the wild type (Fig. S5).

**Fig. 3.**
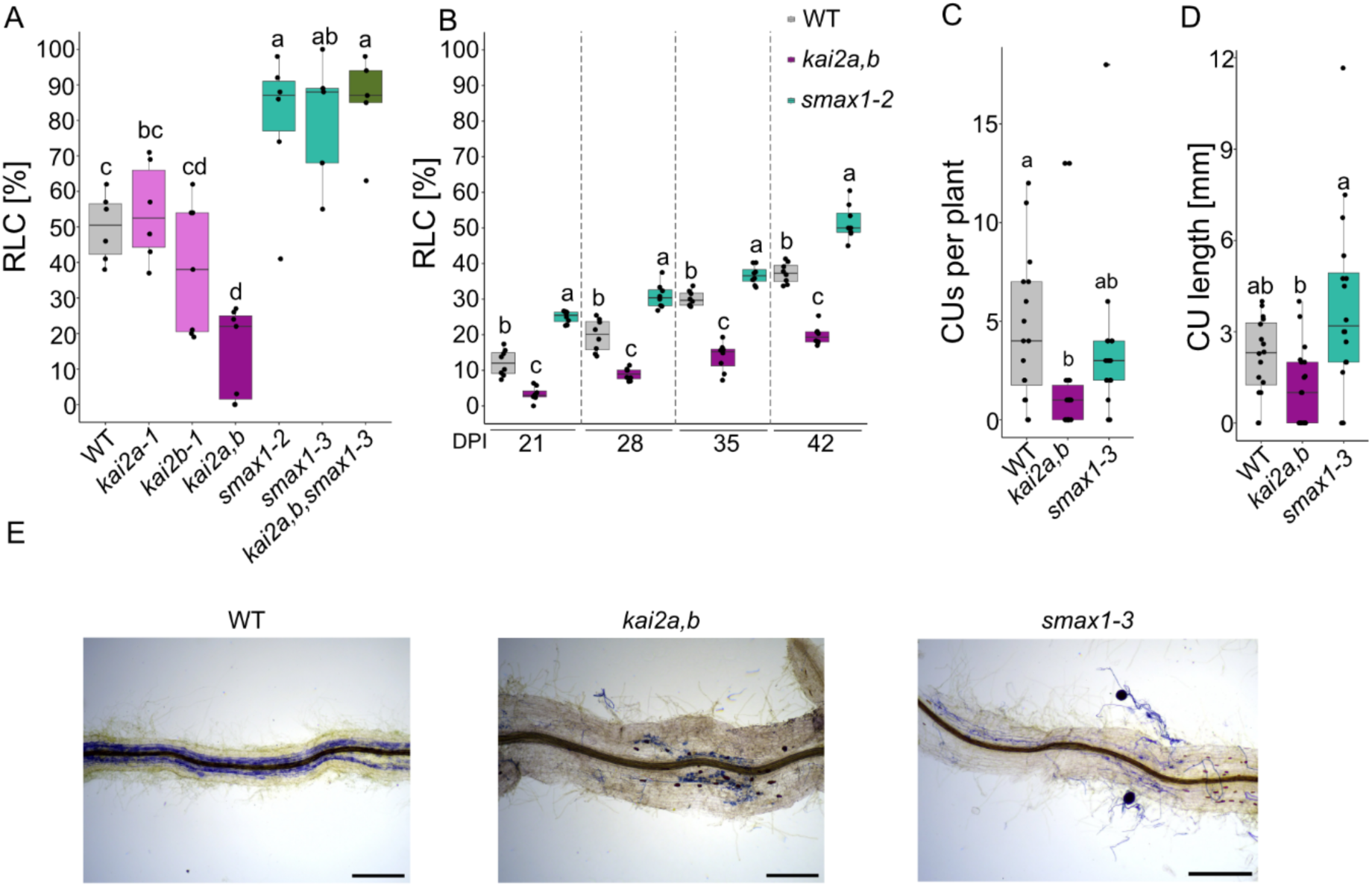
The KAI2-SMAX1 module affects fungal spread in roots of *Lotus japonicus*. Percent total root length colonization (RLC) by *R. irregularis* of the indicated genotypes of *L. japonicus* grown in sand PTC containers, (a) at 28 dpi, (N_plants_ = 6, ANOVA, Tukey, p<0.05, F_6/35_= 16.86); (b) at 21, 28, 35 and 42 dpi (N _plants_ = 8, ANOVA, tukey, F_2/21_ (21 dpi, 28 dpi, 35 dpi, 42 dpi) = 138.6; 80.97; 119.7; 151.5). Different letters indicate different statistical groups. (c) Number of colonization units, (N_plants_ = 16-18, Kruskal-Wallis, Dunn posthoc test, p<0.05, chi-squared = 10.065, df = 2; and (d) average length of colonization units (CU) per plant (N_plants_ = 16-18, Kruskal-Wallis test, with Dunn posthoc test, p<0.05, chi-squared = 12.154, df = 2); in *L. japonicus* wild-type, *kai2a,b* and *smax1* roots grown in sand:clay beads (2:1) after 19 dpi with *R. irregularis*. Experiments in C-D were repeated twice with similar results. Bold horizontal lines represent the median; lower and upper whisker indicate quartile 1 and 4. Lower and upper horizontal lines indicate minima and maxima. Different letters indicate different statistical groups. (e) Bright-field microscopy images of WT, *kai2a,b* or *smax1* roots colonized with *R. irregularis* at 3 wpi with examples of compressed CUs in *kai2a,b*, and elongated hyphae with low arbuscule density in *smax1* CUs. The fungus was stained with acid ink. Scale bars, 500 µM.

### SMAX1_D2_ accumulates in colonized roots

The *L. japonicus* transcriptome data suggested that some genes with increased expression in *smax1* (that may be negatively regulated by SMAX1), can be induced by CO4 treatment in *kai2a,b* roots. These genes may either not be under direct SMAX1 control or SMAX1 is degraded KAI2-independently. To investigate SMAX1 accumulation in *L. japonicus* roots, we used the SMAX1_D2_ degron as a proxy. To this end we introduced an expression cassette encoding the ratiometric reporter *LjUBIQUITIN*_pro_:*LjSMAX1_D2_*-*mScarlet-I-*F2A-Venus-3ΧNLS* (based on the pRATIO3212-SMAX1 ratiometric reporter from Khosla *et al*., 2020) into hairy roots. As the viral *F2A peptide mediates ribosome skipping during translation, two separate, nuclear-localized SMAX1_D2_-mScarlet-I and Venus proteins are produced, and SMAX1_D2_-mScarlet-I mean fluorescence intensity can be used to normalize againg Venus mean fluorescence intensity. Thus, the mScarlet-I/Venus mean fluorescence intensity ratio serves as a proxy for the level of SMAX1 accumulation or degradation. As expected, in non-colonized roots, the mean fluorescence intensity ratio of SMAX1_D2_-mScarlet-I/Venus was higher in nuclei of *kai2a,b* roots, than in wild-type roots, indicating an increased stability of SMAX1_D2_ in absence of KAI2 (Fig. 4c, Table S6). In colonized roots, surprisingly, the SMAX1_D2_-mScarlet-I/Venus ratio was increased compared to non-colonized roots in both wild type and *kai2a,b* (Fig. 4b; Table S6). This indicates that SMAX1 accumulates during colonization or after colonization has occurred and that *Ljkai2a,b* seems to support colonization despite potentially increased accumulation of SMAX1. Furthermore, we observed SMAX1_D2_ accumulation in arbuscule containing cells of both wild type and *kai2a,b* (Fig. 4a-d). This was unexpected as SMAX1 was thought to suppress intracellular colonization. However, due to the asynchronous nature of AM development, colonized roots contain a mix of arbuscules at different developmental stages, which may be associated with different SMAX1 protein levels, and it is possible that SMAX1 accumulation is associated with arbuscule collapse rather than development. Indeed, we found that in wild-type arbuscule-containing cells in nuclear SMAX1_D2_-mScarlet-I/Venus ratios ranged from below to above one (Fig. 4a-b, d), suggesting that degradation of SMAX1_D2_ may differ among different stages of arbuscule development. However, in *kai2a,b* roots, we only saw nuclei in arbuscule-containing cells with SMAX1_D2_-mScarlet-I/Venus ratios of above 3, indicating that SMAX1 degradation in arbuscule-containing cells may be dependent on presence of KAI2 (Fig. 4a-b, d). To understand how SMAX1_D2_ stability may differ across root tissue-types we assigned the nuclei from mock and AM inoculated roots in Fig. 4c to their respective cell-type (Fig. 4d). Both under mock and AM conditions SMAX1_D2_ accumulation in the epidermis was similar in roots of wild type and *kai2a,b* (Fig. 4d). However, in the wild type under both mock and AM conditions, SMAX1_D2_ levels in the cortex and vasculature were lower than in the epidermis (Fig. 4d). In *kai2a,b*, SMAX1_D2_ levels were similar in epidermis, cortex and vasculature in mock conditions, whereas in AM roots SMAX1_D2_ levels were higher in the cortex and vasculature than in the epidermis (Fig. 4d). An increased SMAX1 accumulation in cortex and vasculature, and the increased colonization in AM roots may contribute to the decreased RLC in *kai2a,b*, as compared to wild-type RLC. In non-colonized *kai2a,b* roots we found a few cells with SMAX1_D2_-mScarlet-I/Venus ratios below 1, indicating that SMAX1_D2_ can be degraded KAI2-independently, at least at low level and in a few cells. However, as in *kai2a, b* mutants the SMAX1_D2_-mScarlet-I/Venus ratios were generally above 1, we do not have convincing evidence for KAI2-independent SMAX1 degradation being responsible for colonization of *kai2a,b* mutants, except if the difference in SMAX1_D2_-accumulation between mock and AM roots would indicate increased SMAX1_D2_ degradation in mock roots. Instead, the data rather suggest that colonization of *kai2a, b*, may occur despite the accumulation of SMAX1 and that SMAX1 accumulation in AM roots could potentially be caused by a stronger stabilization of SMAX1_D2_ as compared to Venus in AM conditions, or a specific increase in SMAX1_D2_ translation.

**Fig. 4.**
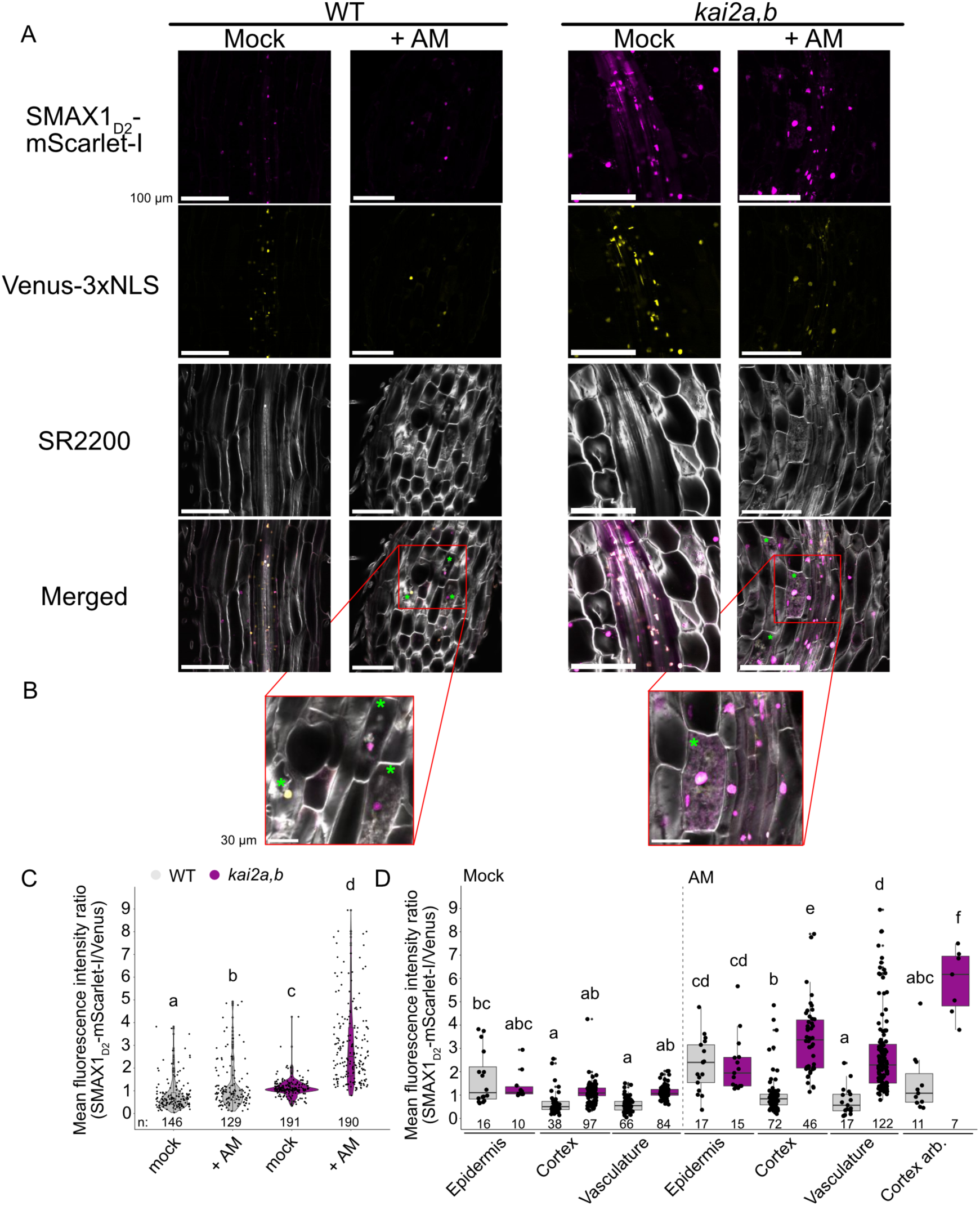
SMAX1_D2_-mScarlet-I accumulates in colonized roots. (a) Representative confocal images of vibratome sectioned (100 µm) *L. japonicus* WT and *kai2a,b* hairy roots colonized (AM) or not colonized (Mock) by *R. irregularis* and expressing *UBI_pro_:SMAX1_D2_-mScarlet-I-*F2A-Venus3xNLS*; scale bars = 100 µm. (b) Close up of arbuscule containing cells from images in (a); scale bars = 30 µm. Green asterisks indicate arbuscule-containing cells. Note the yellow nucleus (indicating absence of SMAX1) in a WT arbuscule-containing cell. (c) Ratios of mean intensities of nuclear SMAX1_D2_-mScarlet-I signal to nuclear Venus signal measured in confocal microscopy images of hairy root longitudinal sections as shown in a. (d) SMAX1_D2_-mScarlet-I/Venus ratio per root tissue type from mock or AM inoculated WT and *kai2a,b* roots. Different letters indicate statistically different groups. For (c) and (d) SMAX1_D2_-mScarlet-I/Venus ratios were analyzed using a linear mixed-effects model on log-transformed data, including log(Venus) as a covariate. Estimated marginal means were compared between groups using Tukey-adjusted post hoc tests. Different letters indicate statistically significant differences (p < 0.05). In D, bold horizontal lines represent median value; lower and upper whisker indicate quartile 1 and quartile 4. Lower and upper horizontal lines indicate minima and maxima. For c, the experiment was repeated twice with similar results. Pictures for ratio analysis were taken from at least 3 separate roots per condition, every root constituting a separate transformation event. Larger points indicate data point from 1 nucleus, smaller dots indicate potential statistical outlier.

## Discussion

Here we show that in the dicotyledons *L. japonicus, P. sativum* and *N. benthamiana*, KAI2-signalling contributes quantitatively to AM development, but is not essential for root colonization. This means that the requirement of KAI2-signalling for AM symbiosis varies across angiosperms, as an absence of colonization in rice *d14l/kai2* roots and the strong reduction described for roots of *Bdkai2* indicate a more critical role in these species (Gutjahr *et al*., 2015; Meng *et al*., 2022). Any of a plethora of (developmental and cell wall) traits that differ between the roots of these poaceous grasses and those of the dicotyledon species investigated in this study may affect the role of karrikin signalling in AM symbiosis. It is, therefore, likely that this divergent requirement across angiosperms cannot be explained by a single gene or trait. Moreover, the difference in the contribution of KAI2-signalling to AM does not seem to align with a monocotyledon-dicotyledon divide, as in the monocotyledon barley KAI2 also has only a quantitative impact on AM.

We found that surprisingly mutation of KAI2 has a stronger impact on the transcriptome of non-stimulated *L. japonicus* than on *B. distachyon* roots, including several genes known to be required for AM. Thus, apart from differing among the species, gene expression analysis did not provide a clear lead on why KAI2 has a stronger impact on AM in *B. distachyon* than in *L. japonicus*. However, other differentially expressed genes with yet unknown function in AM may also explain these differences. In addition, transcriptional responses to GSE and CO4 and the impact of KAI2 on these responses largely differ between *L. japonicus* and *B. distachyon,* suggesting that receptors for fungal molecules as well as karrikin signaling pathway are differentially wired in the two species or potentially affected by the interaction of karrikin signaling with other hormone signalling pathways. However, we show that in *L. japonicus*, genes crucial for AM symbiosis, such as *CCaMK*, can be induced in *kai2a,b* roots slightly by CO4 treatment and significantly by root colonization. This effect of CO4 treatment was not observed for *B. distachyon* roots and this unresponsiveness may contribute to the stronger AM phenotype. Some of the genes induced by CO4 in *Ljkai2a,*b roots also show increased expression in *smax1* roots, suggesting they may be direct or indirect SMAX1-targets. It is possible that alternative and redundant pathways regulate these genes, and are able to override the transcriptional repression due to SMAX1 accumulation. Alternatively, SMAX1 could be targeted for degradation through mechanisms acting in parallel with the karrikin receptor complex.

We used SMAX1_D2_-mScarlet-I as a marker for SMAX1 degradation (Khosla *et al*., 2020) and observed increased accumulation of SMAX1_D2_-mScarlet-I in *kai2a,b* roots as compared to wild type, confirming that KAI2 is required for SMAX1 degradation. However, in some cells of *kai2a,b* roots we found a relative mScarlet-I fluorescence intensity lower than 1, indicating that SMAX1 can be degraded locally to a small degree in *Ljkai2a,b* roots, although less efficiently than in wild type and only in few cells. It is thus tempting to speculate that SMAX1 degradation, by proxy of SMAX1_D2_ degradation, can occur independently of KAI2. KAI2-independent degradation of SMAX1 has been shown in *Arabidopsis* during osmotic stress and to depend on the strigolactone receptor D14 stimulated treatment with the synthetic strigolactone analog GR24^5DS^ (Li *et al*., 2022a). It will be interesting to determine whether D14-mediated SMAX1 degradation also occurs in *L. japonicus* roots or whether other mechanisms are at play. In non-colonized wild-type roots we observed differences in mScarlet-I/Venus ratio across tissue types, with lower levels of SMAX1_D2_-mScarlet-I accumulation in the cortex and vasculature than in the epidermis. This suggests differences in activation of karrikin signalling across tissue types in *L. japonicus*, especially because these tissue-specific differences were not observed in *kai2a,b,* where the median value was close to 1 across all tissues.

Upon root colonization, we observed a surprising increase in the accumulation of SMAX1_D2_-mScarlet-I in roots of both wild type and *kai2a,b*, as compared to non-colonized roots. This was particularly pronounced in *kai2a,b*, especially in arbuscule-containing cells and suggests stabilization of SMAX1 in these cells causing SMAX1 to be more stable than Venus. Alternatively, SMAX1 translation could be preferred over Venus translation in these cells due to an unknown mechanism. Based on the rice *d14l*/*kai2* phenotype (Gutjahr *et al*., 2015; Meng *et al*., 2022) it may have been suggested that removal of SMAX1 is required for fungal into plant cells. It is therefore, at first sight unexpected to see SMAX1_D2_ accumulation in arbuscule-containing cell. However, the arbuscule life-cycle undergoes different stages from formation to degeneration and collapse (Gutjahr & Parniske, 2013) and SMAX1 accumulation could be associated with the initiation of arbuscule collapse, for example due to a reduction in KAI2 expression and signalling due to accumulating phosphate in the arbuscule-containing cell (Villaécija-Aguilar *et al*., 2022) or because other hormone pathways are activated in arbuscule-containing cells such as ethylene signalling, which was recently shown to promote SMAX1 accumulation, either by increased synthesis or decreased degradation (Das *et al*., 2025).

The observed stabilization of SMAX1 during AM could be important for limiting the fungal spread through the root at a later stage of the symbiosis. In our study we present evidence that karrikin signalling modulates the number size, and morphology of colonisation units. *Ljkai2a,b* roots, in which SMAX1 and also SMAX1_D2_ accumulate more strongly than in wild type, contain fewer and shorter CUs. Also in presence of ethylene, which boosts accumulation of SMAX1, colonization units are short and compact (Das *et al*., 2025), resembling the *kai2a,b* phenotype. In addition, we found that *kai2a,b* mutants accumulate higher levels of SMAX1_D2_ in the vasculature and cortex. It has recently been shown in *M. truncatula* that the GRAS protein DELLA needs to move from the vasculature to the cortex to enable arbuscule development (An *et al*., 2025). It is possible that SMAX1 in both tissues reduces arbuscule development by sequestering DELLA (Xu *et al*., 2023; Hong *et al*., 2026).

As karrikin signalling has been shown to affect strigolactone biosynthesis genes, and rice *smax1* mutants exhibit increased strigolactone exudation, the reduced number of CUs in *kai2a,b* may also be explained by reduced strigolactone biosynthesis (Choi *et al*., 2020). However, this may not be the only reason, as *smax1* did not show an increase in CUs.

Taken together, as we observed only minor evidence for SMAX1_D2_ degradation in *Ljkai2a,b* mutants we propose that either a modification of SMAX1 activity (e. g. by posttranslational modification), rendering it less active despite its presence, or induction of AM-relevant gene expression by redundant pathways that may bypass accumulated SMAX1, may be the cause for the reduced requirement of KAI2-signalling in AM symbiosis in *L. japonicus* as compared to *B. distachyon* or rice. This redundancy may result from the combined activity of multiple regulators targeting genes important for AM symbiosis, as well as from additional pathways regulating the stability or activity of SMAX1. Future work aimed at identifying these players will be important for understanding the mechanistic basis of the variation across plant species in KAI2 requirement for AM symbiosis.

## Acknowledgements

We thank Alexandre de Saint Germain (INRA Versailles, France) for pea mutant seeds. We are grateful to Andrea Müller (IPK Gatersleben) for the technical assistance for the generation of the *N. benthamiana kai2* quadruple mutant plants and Antje Bolze (MPI of Molecular Plant Physiology) for genotyping the *N. benthamiana kai2* quadruple mutant lines. We thank Dr. Christine Wurmser (NGS@TUM) for the preparation of RNA-sequencing libraries. This study was supported by the Deutsche Forschungsgemeinschaft (DFG), first by their Emmy Noether program (259604726) to CG and then by the transregio collaborative research center TRR356 ‘Plant-Microbe’ (491090170) to CG (project A02) and NK (project Z03) and by a core grant from the Max Planck Society to CG.

## Author contributions

KB, ST and CG conceived the study. KB, ST, SC, KV and PC conducted phenotypic analyses. ST, SC and KV performed mutant complementation analyses. GH generated *N. benthamiana kai2* quadruple mutants and ST identified and characterized the mutations (with help from Antje Bolze) and characterized the mutants. ST established the conditions for and performed the RNAseq experiment. AF, MM NK, ST and KB analysed the RNAseq data. KB analysed SMAX1_D2_ accumulation. KB generated the final figures and wrote the first draft of the manuscript. CG edited the manuscript and supervised the study. NK and CG acquired funding.

---

**Fig. S1.**
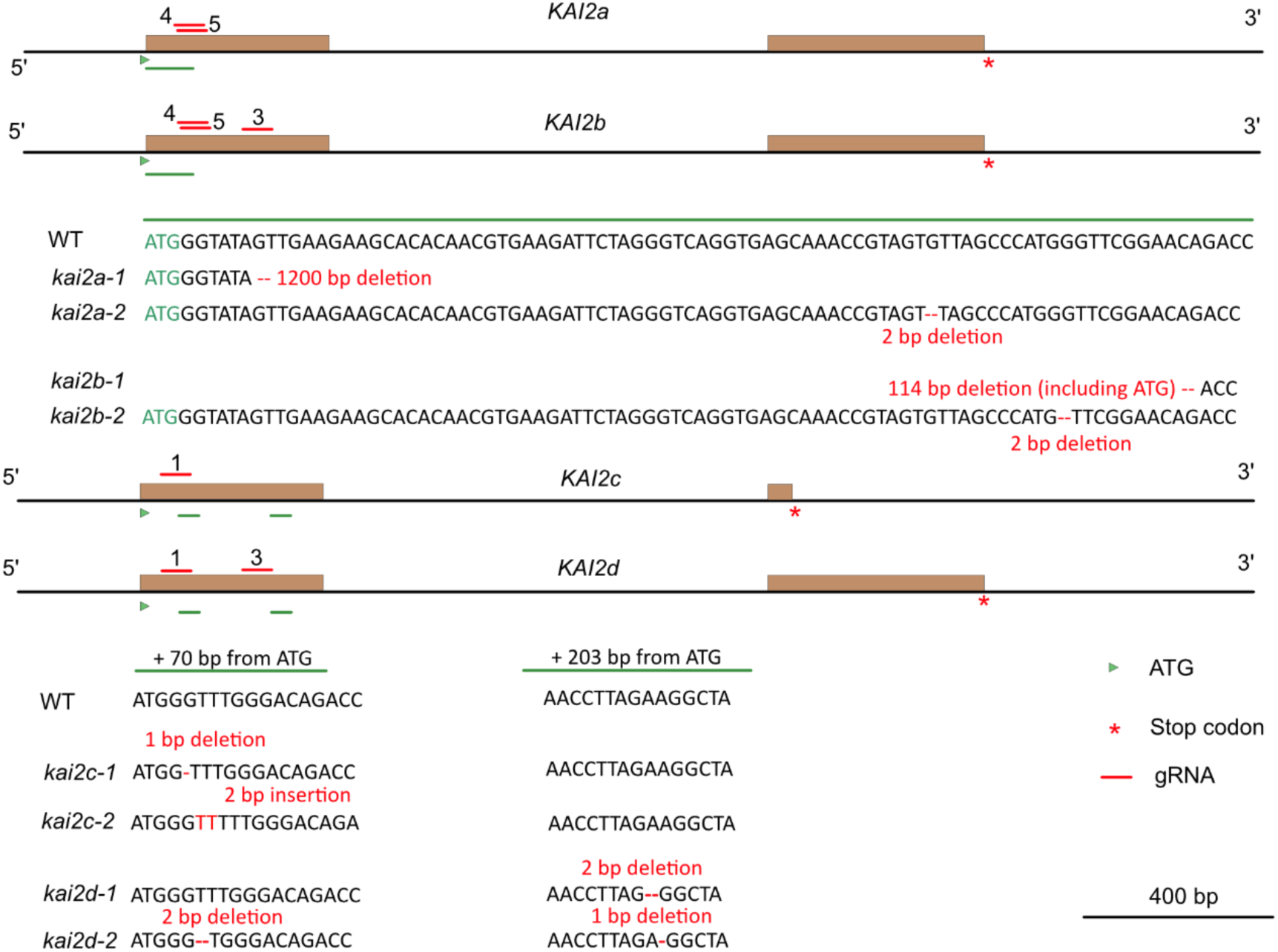
Schematic representation of cas9-mediated mutations in *kai2* paralogs in *N. benthamiana*. Schematic representation of the four conserved *KAI2c* homologs in *N. benthamiana* and their edits resulting in frame shifts or large deletions. Brown boxes indicate exons areas on black lines between brown boxes indicate introns and 5’ or 3’ UTRs. Green bars represent gene edits. Exon size is based on to the *SolGenomics* genome browser of *N. benthamiana* genome version v1.0.1.

**Fig. S2.**
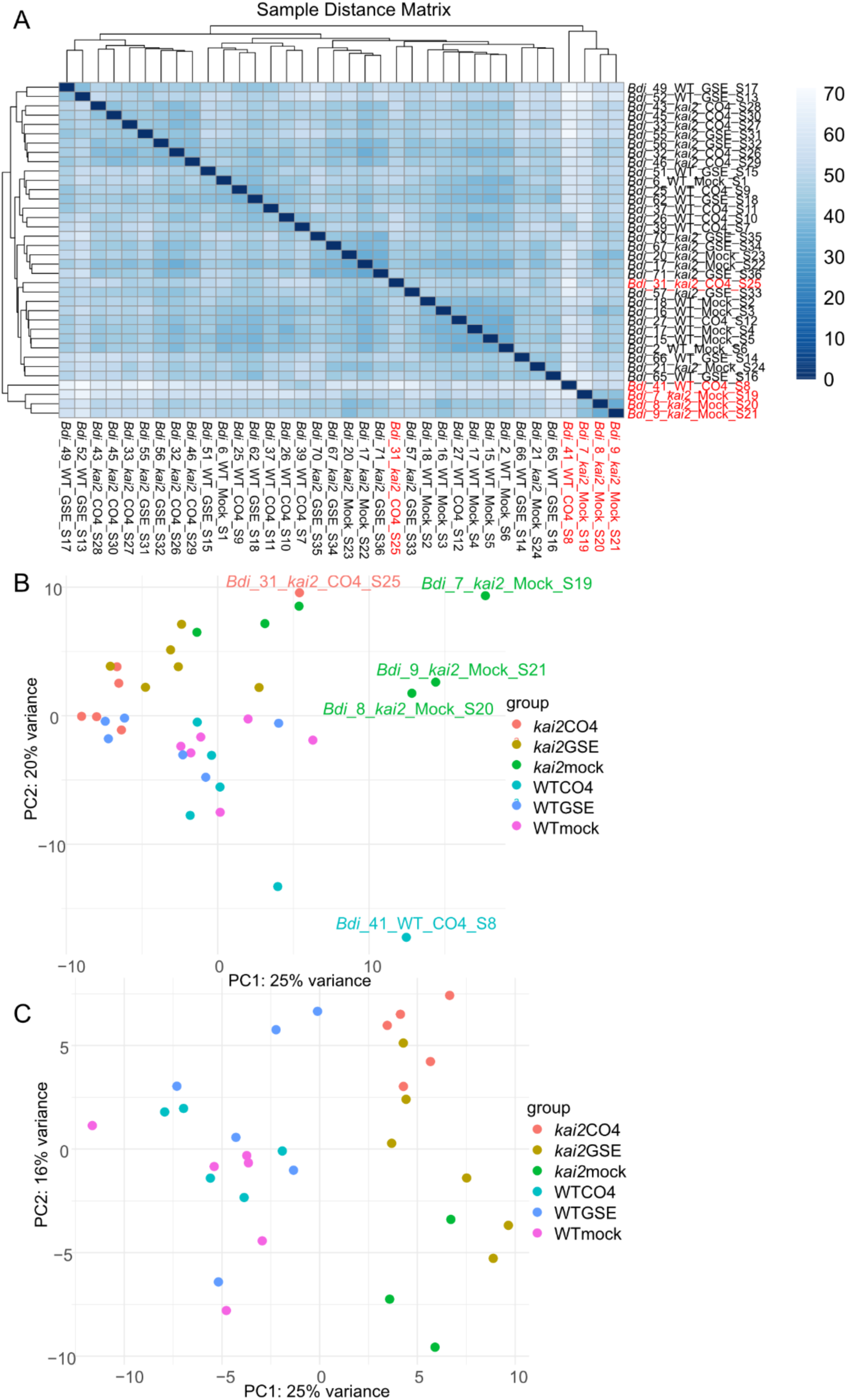
Sample distance hierarchical clustering and PCA of *B. distachyon* transcriptome. (a) Hierarchical clustering of the sample distance between all *B. distachyon* root transcriptome samples. Red labels indicate samples that were removed from the final analysis because they were too distant from their cluster in the Principal Component Analysis (PCA) (b) PCA of *B. distachyon* transcriptome samples. Samples with a name label were removed from further analysis. (c) PCA *of B. distachyon* root transcriptome after removal of samples marked in (a-b).

**Fig. S3.**
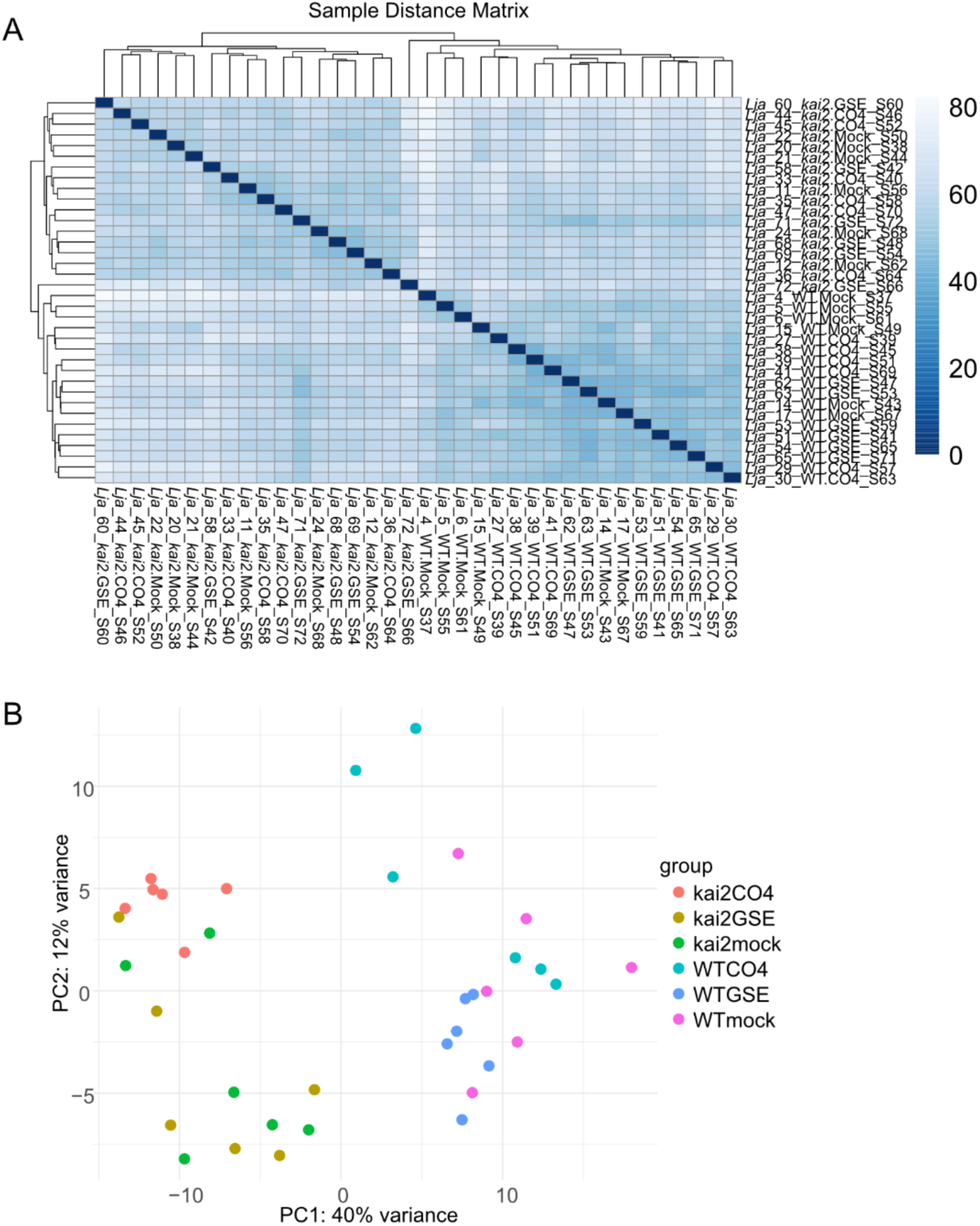
Sample distance hierarchical clustering and PCA of *L. japonicus* transcriptome. (a) Hierarchical clustering of the sample distance between all *L. japonicus* transcriptome samples. (b) Principal Component Analysis (PCA) of *L. japonicus* transcriptome samples.

**Fig. S4.**
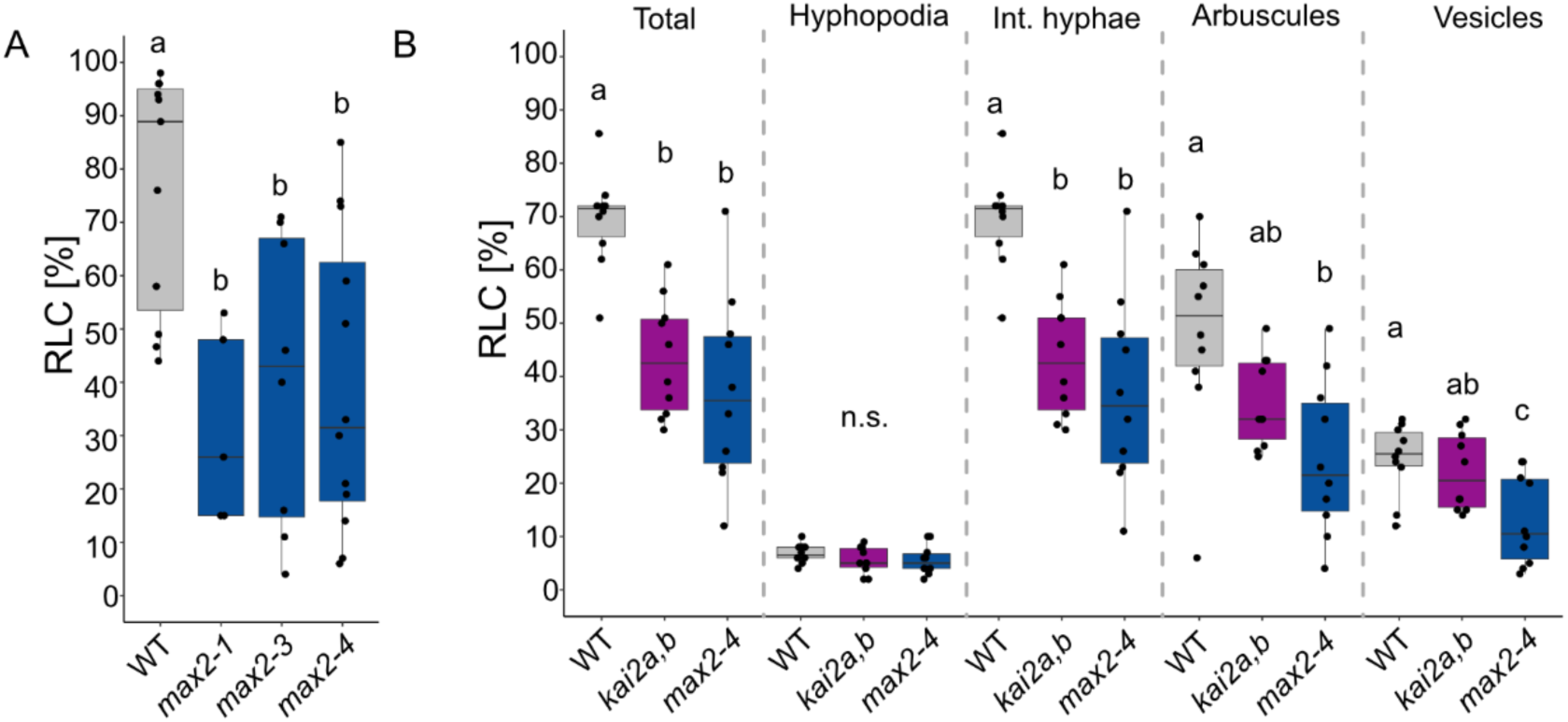
Reduction of total root length colonization in karrikin signalling mutants. (a) Percent total root length colonization (RLC) in *L. japonicus* wild type, *max2-1, max2-3,* and *max2-4* grown in sand vermiculite open pot cultures for 42 days post inoculation (dpi). ANOVA, Tukey, p<0.05. F_3/32_=6.04. Different letters indicate statistically significant different groups. (b) Percent RLC for fungal colonisation structures in wild type, *kai2a,b* and *max2-4* roots from Fig. 1a. ANOVA, Tukey, p<0.05. F_2/27_(Total/Int. hyphae/ Arbuscules/Vesicles)=17.06/6.951/17.19/6.534.

**Fig. S5.**
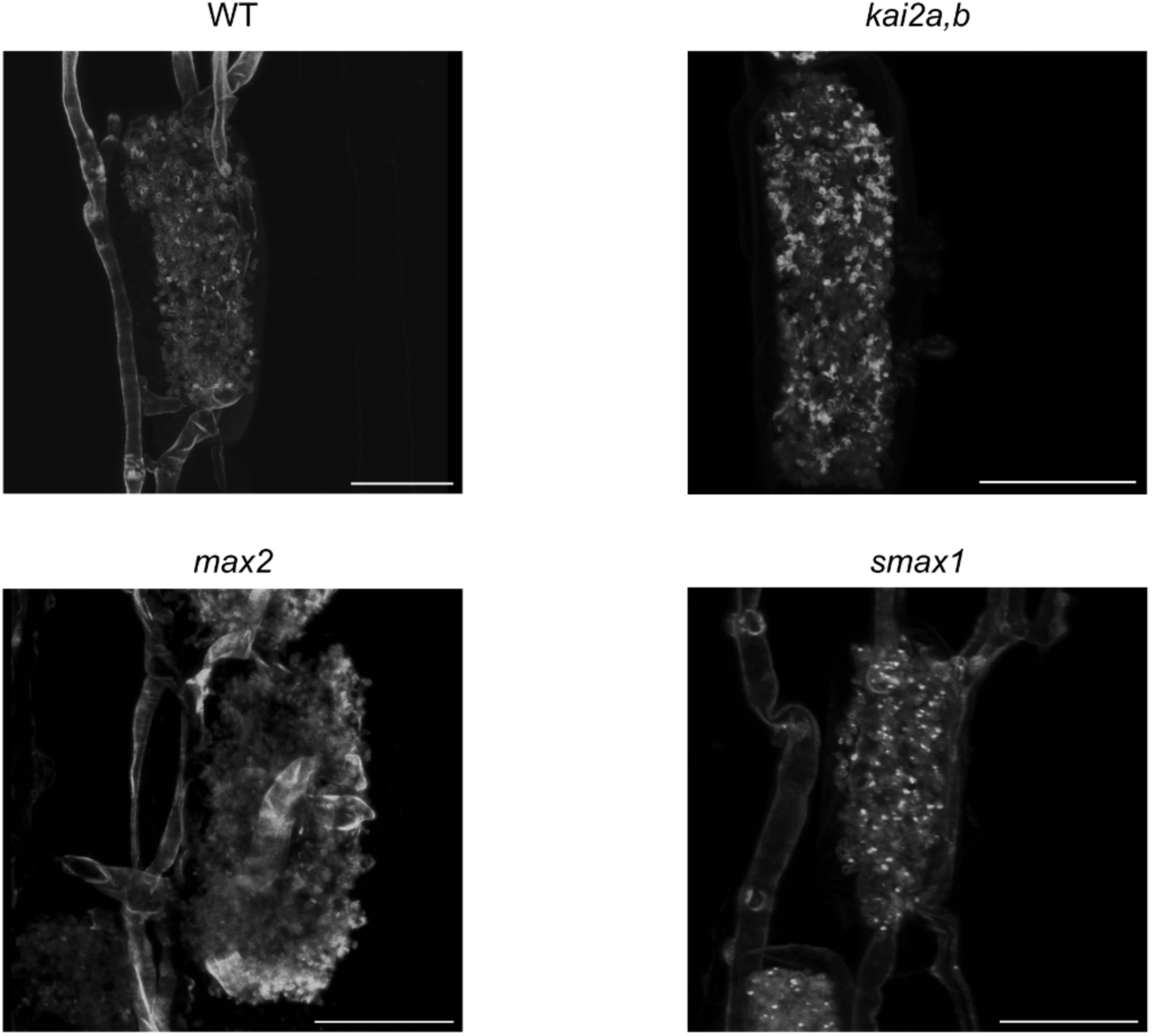
Confocal microscopy images of representative arbuscules in wild-type, *kai2a,b, max2,* and *smax1* root cortex cells. Roots were grown in open pot sand cultures for 21 days with spores of *R. irregularis* were. The fungus was stained with Wheat Germ Agglutining(WGA)-AlexaFluor-488. Scale bars = 25 µm.

**Fig. S6.**
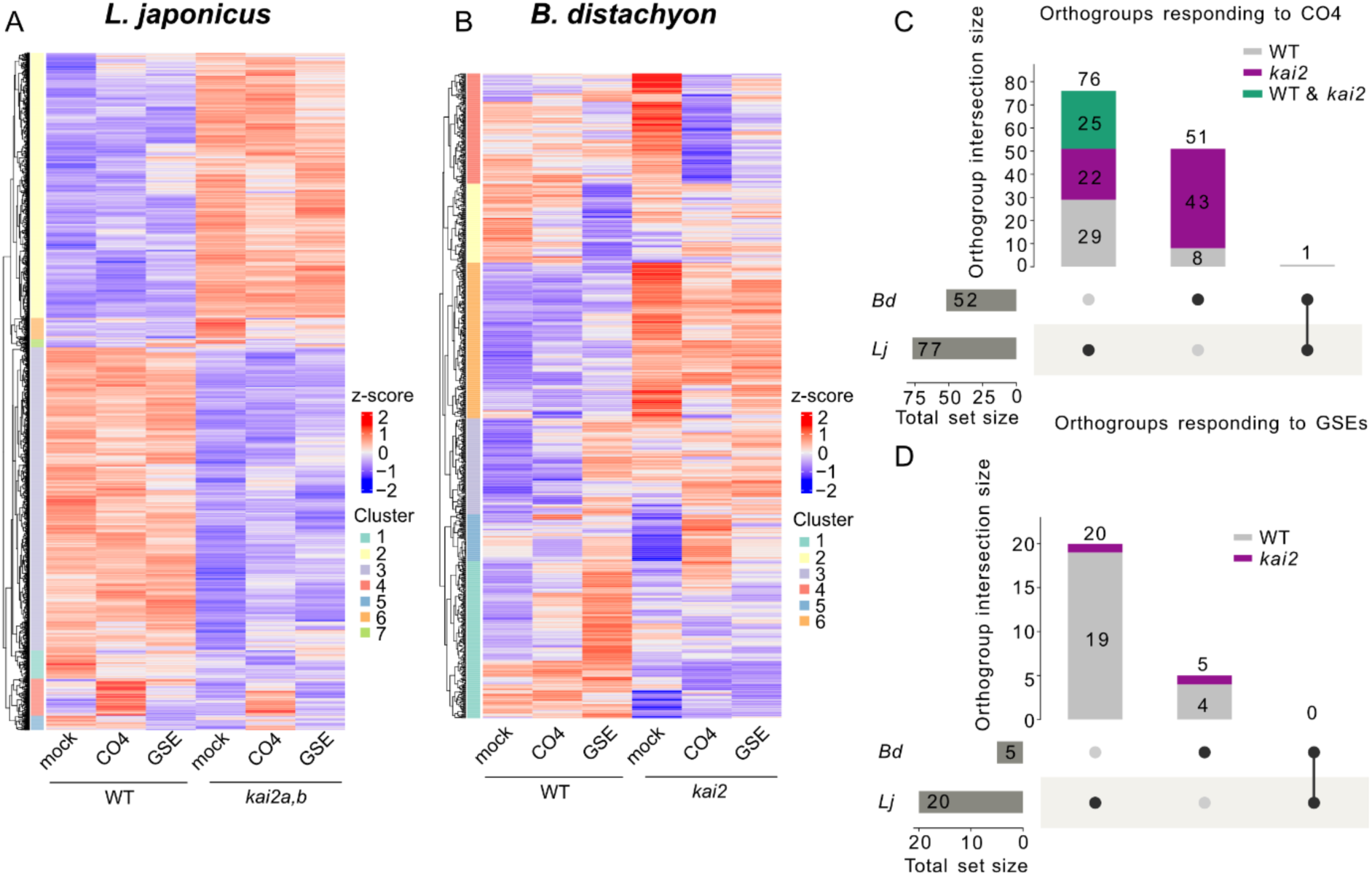
Transcriptomes of *L. japonicus* and *B. distachyon* wild type or *kai2a,b/kai2* roots responding to CO4 or GSE treatment. (a-b) K-means clustered heatmaps visualizing the relative expression value (z-score) of all DEGs (padj<0.05) across the transcriptomes of (a) *L. japonicus* (2537 DEGs, k=7) or (b) *B. distachyon* (809 DEGs, k=6). (c-d) Upset plots showing shared orthogroups of the (c) CO4 or (d) GSE responsive DEGs in *L. japonicus* (*Lj*) and *B. distachyon* (*Bd*). Set size indicates the total amount of orthogroups that respond to CO4 or GSE treatment in either *L. japonicus* or *B. distachyon,* and intersection size indicates number of responsive orthogroups either unique to *L. japonicus* or *B. distachyon*, or shared across the species. Colour indicates whether genes in the orthogroups respond to treatment in wild type (gray), *kai2* (purple) or both (green).

**Fig. S7.**
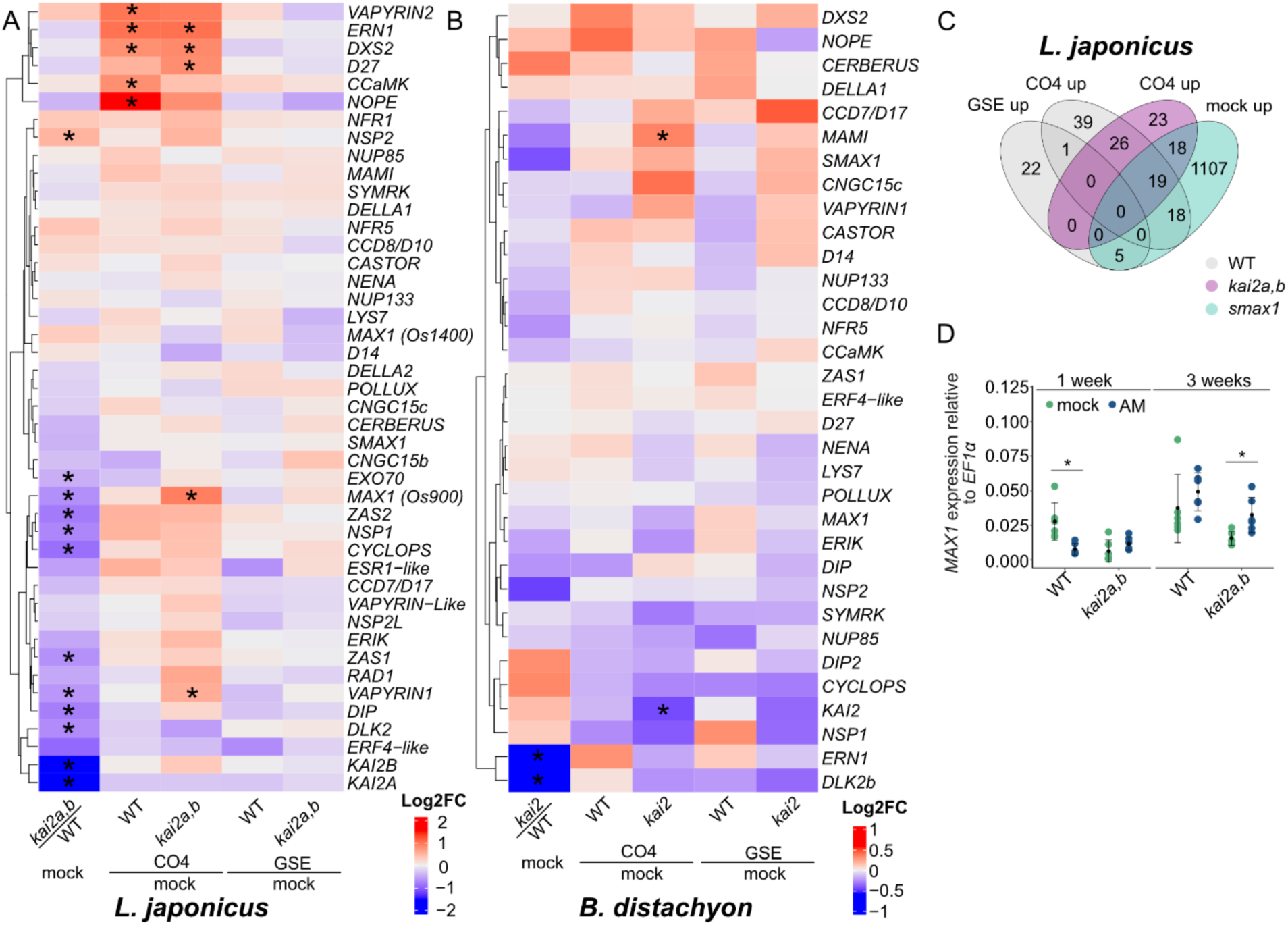
Transcriptional response of AM-, karrikin signalling-, and strigolactone biosynthesis and signalling genes to CO4 or GSE treatments in *L. japonicus* and *B. distachyon* roots. Heatmaps visualising Log2FC of genes important for AM symbiosis, or involved in karrikin or strigolactone signalling in the (a) *L. japonicus* or (b) *B. distachyon* root transcriptome. Asterisks indicate statistically significant fold change (padj<0.05). (c) Venn diagram showing overlap in number of genes significantly upregulated (Log2FC>0; padj<0.05) in wild-type + GSE or CO4, *kai2a,b* + CO4 and *smax1* at mock condition in *L. japonicus*. (d) Expression of *MAX1* relative to *LjEF1α* in roots of WT or *kai2a,b*, at 1 or 3 wpi without (mock) or with spores of *R. irregularis* (AM). (ANOVA, estimated marginal means contrast for AM treatment, p<0.05; (F_1/40_ = 4.313, * p<0.05). Black dot indicates mean value, black error bars represent standard deviation of the mean.

**Fig. S8.**
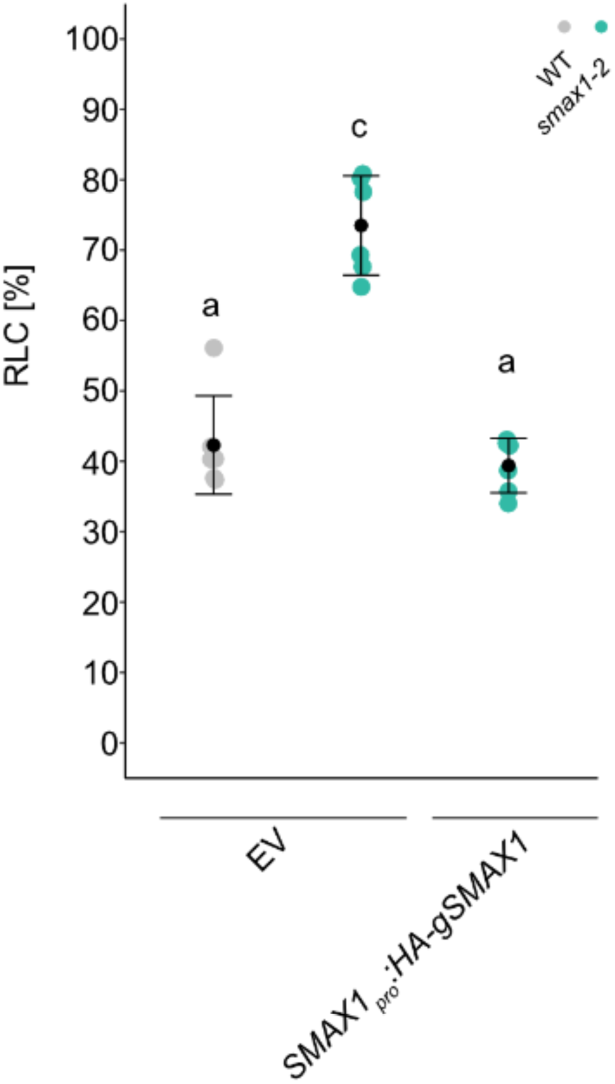
Complementation of *Ljsmax1* with *gSMAX1* leads to reduced colonization. Percent total root length colonization (RLC) of *L. japonicus* wild type and *smax1-2* mutants transgenic roots after *Agrobacterium rhizogenes*-mediated transformation with empty vector (EV) or *SMAX1_pro_:HA-gSMAX1* at 28 days post inoculation with *R. irregularis*. Plants were grown in open pot sand cultures. Different letters indicate different statistical groups. ANOVA, Tukey, p<0.001. F_2/15_=56.6. Black dot indicates mean value, black error bars represent standard deviation of the mean value.

## Supplementary Tables

**Supplementary Table 1.**
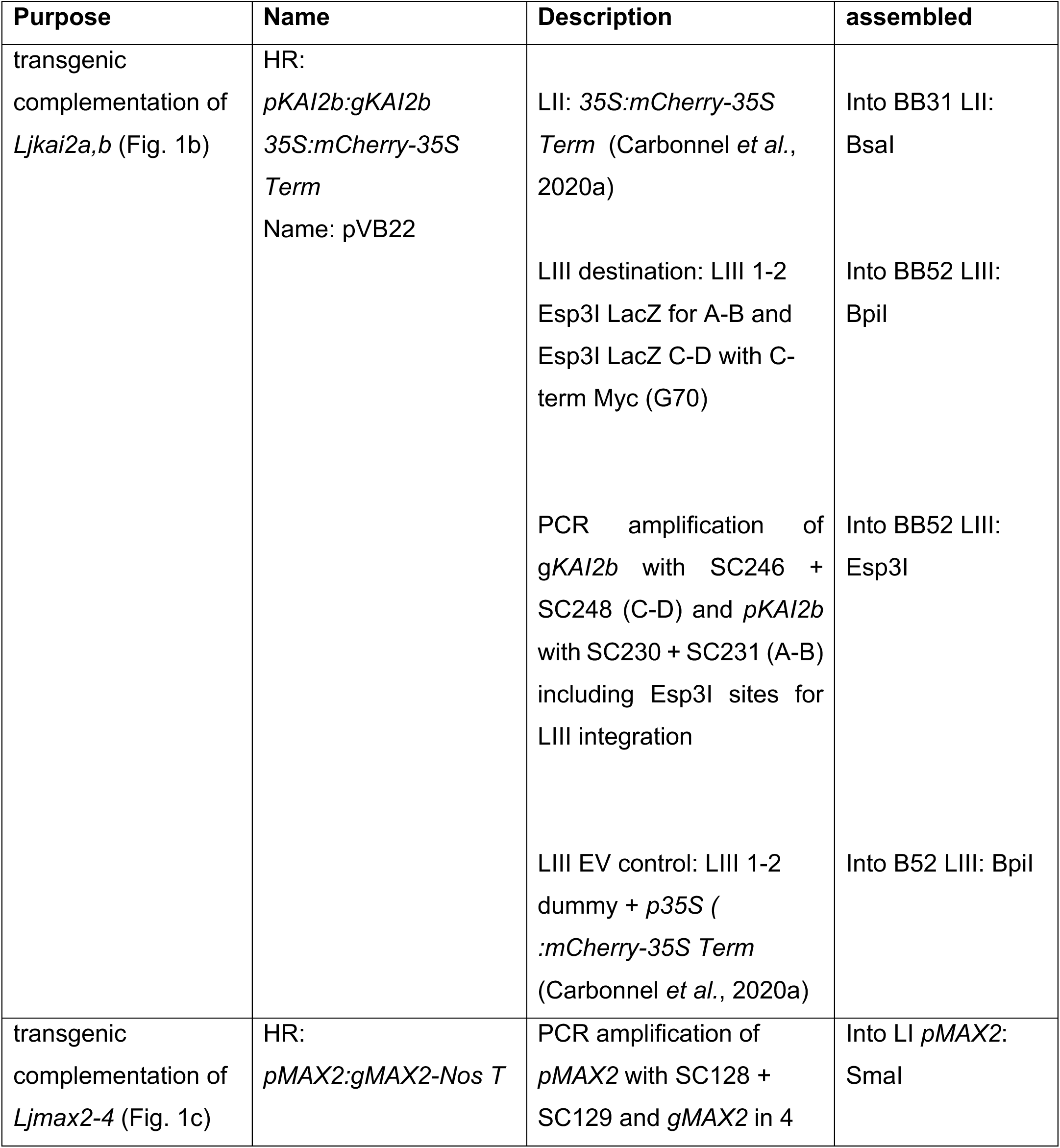
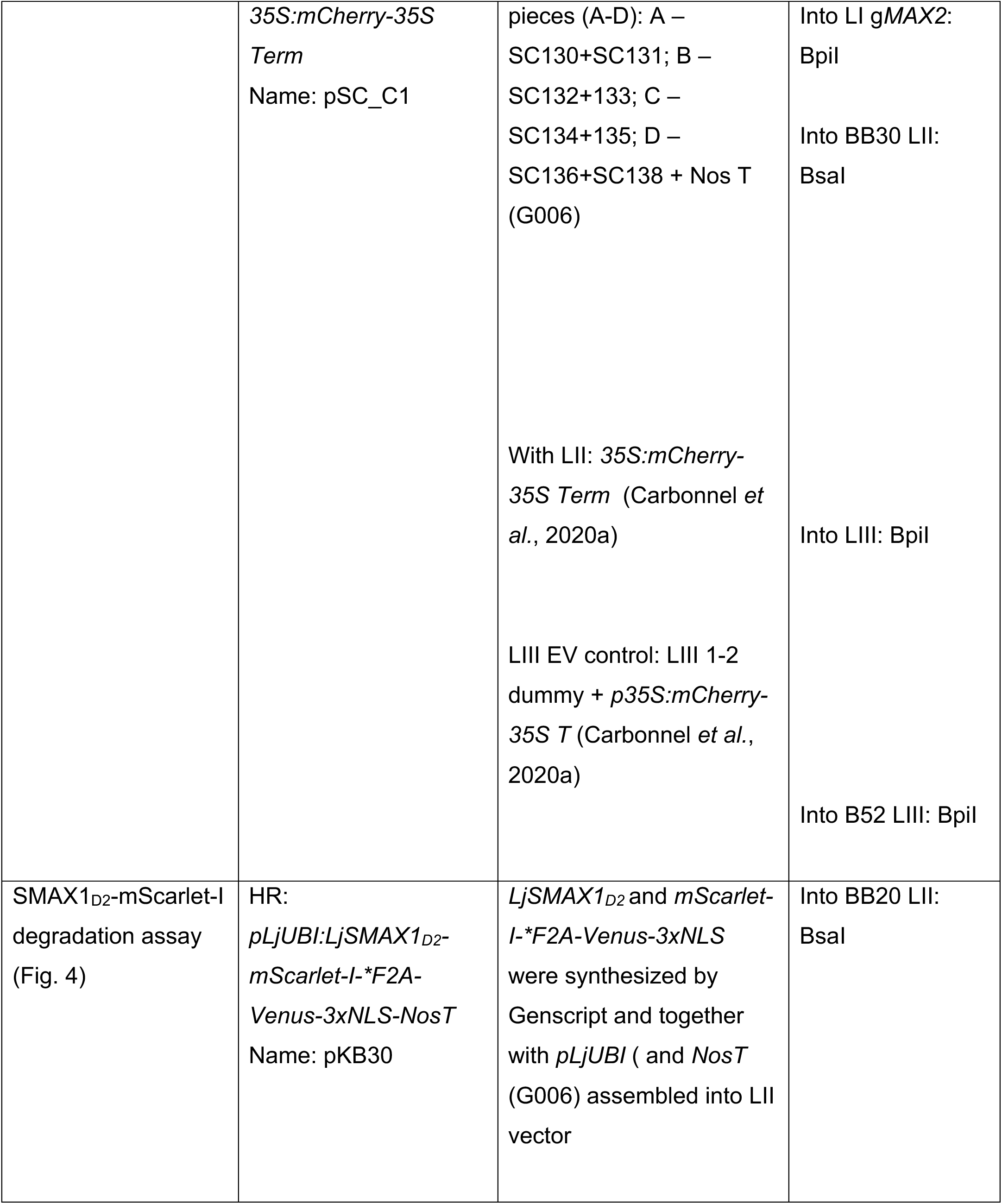
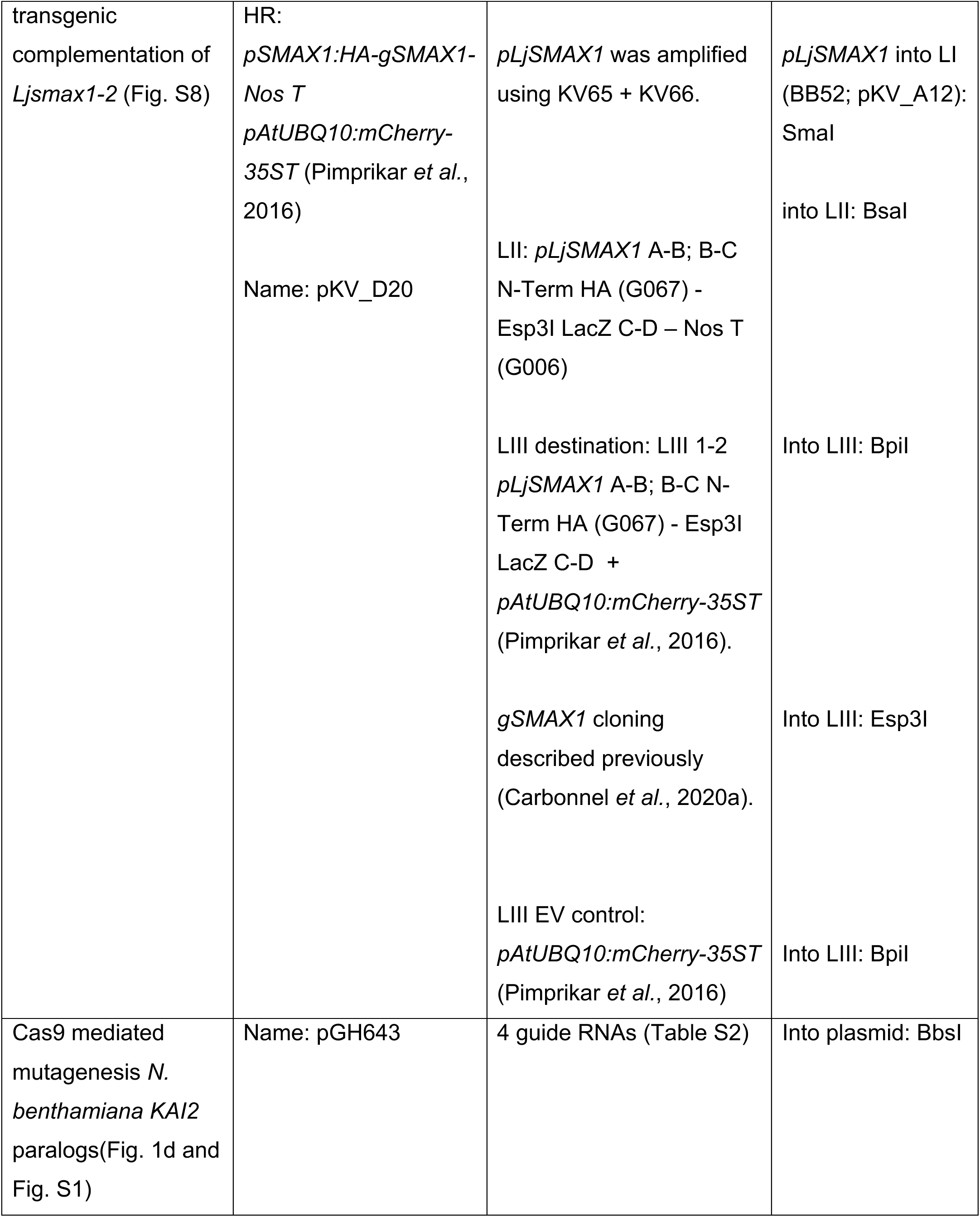
Plasmids used in this study All plasmids for this study were produced using the Golden Gate toolbox (Binder *et al*., 2014), except pGH643. EV: Empty Vector. HR: Hairy root transformation.

**Supplementary Table 2.**
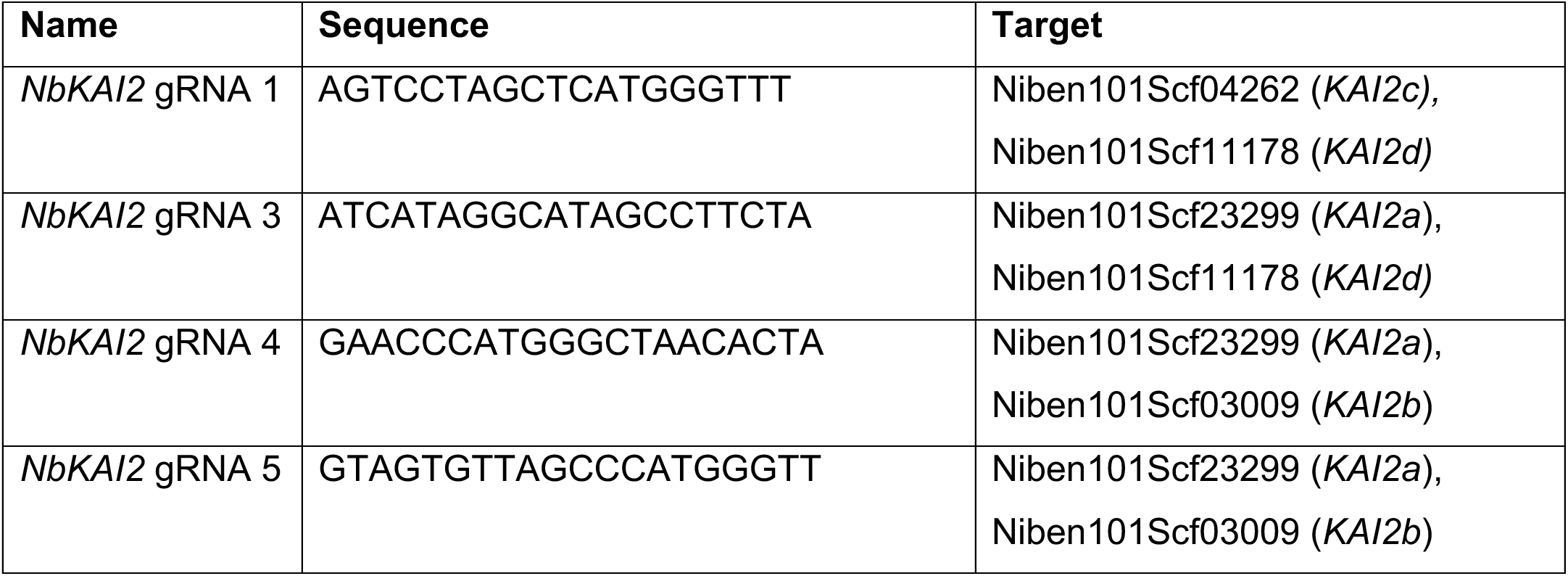
Sequence of gRNAs used for *NbKAI2a, b,c,d* editing.

**Supplementary Table 3.**
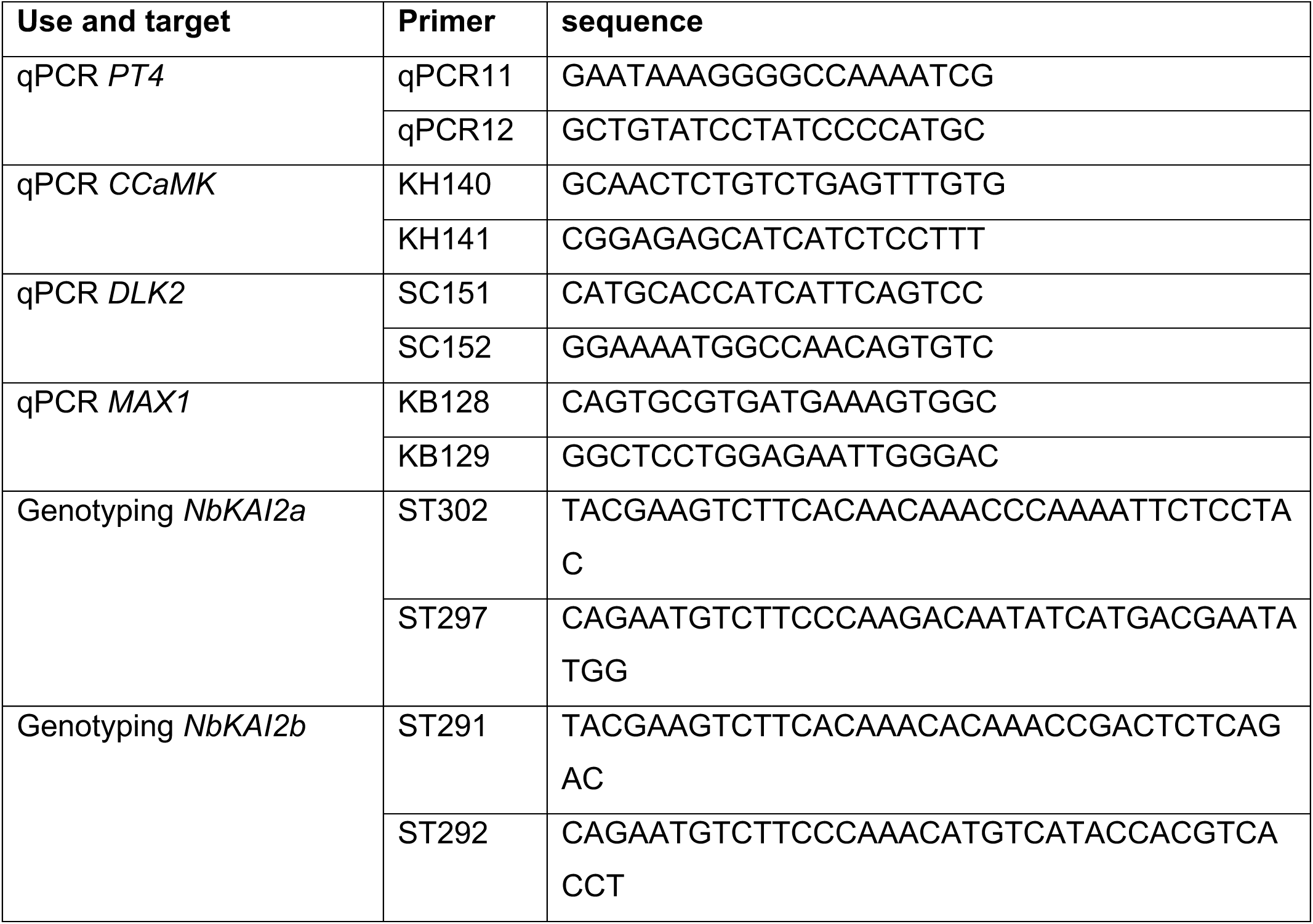
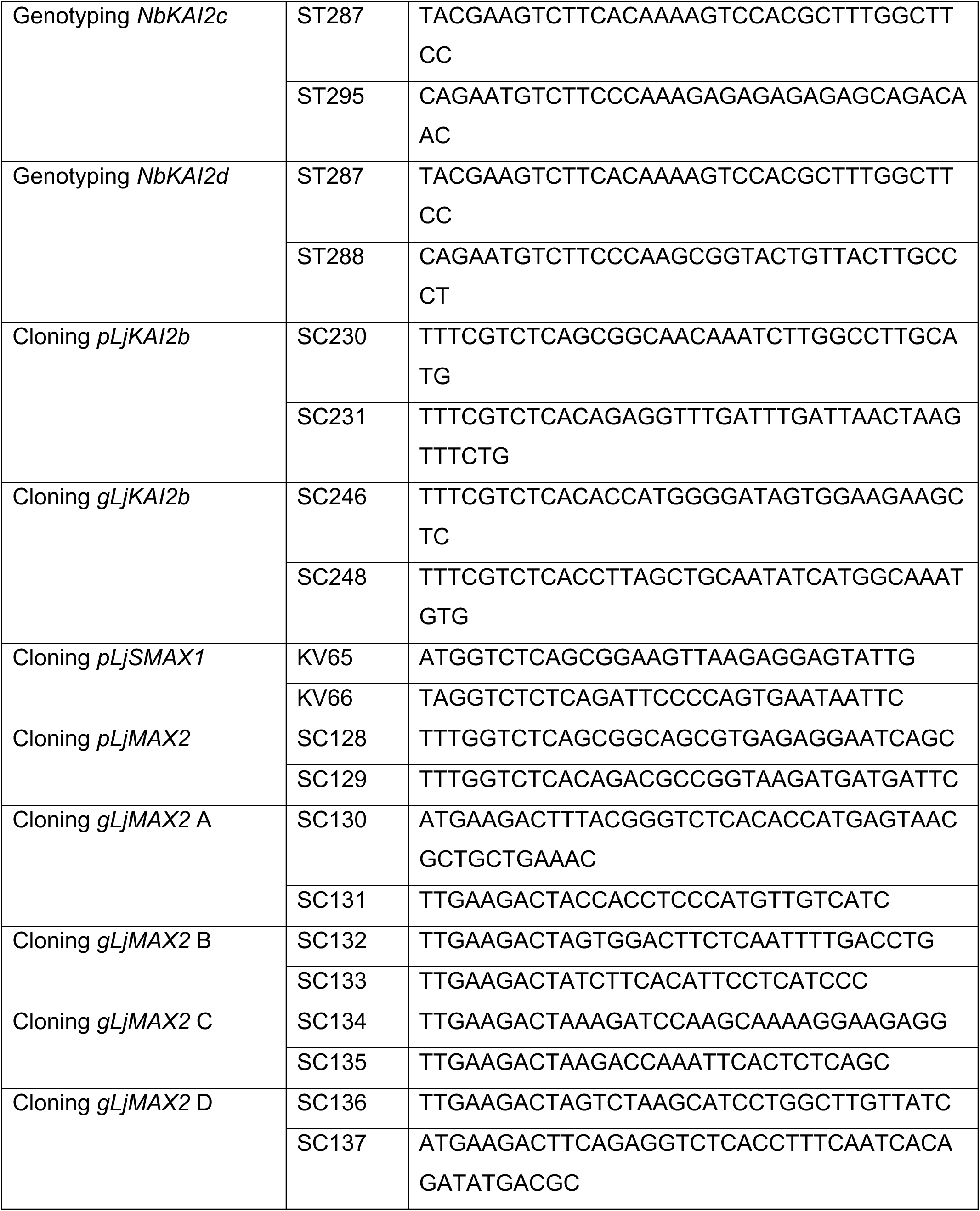
Primers used for qPCR, genotyping and cloning.

**Supplementary Table 4.**
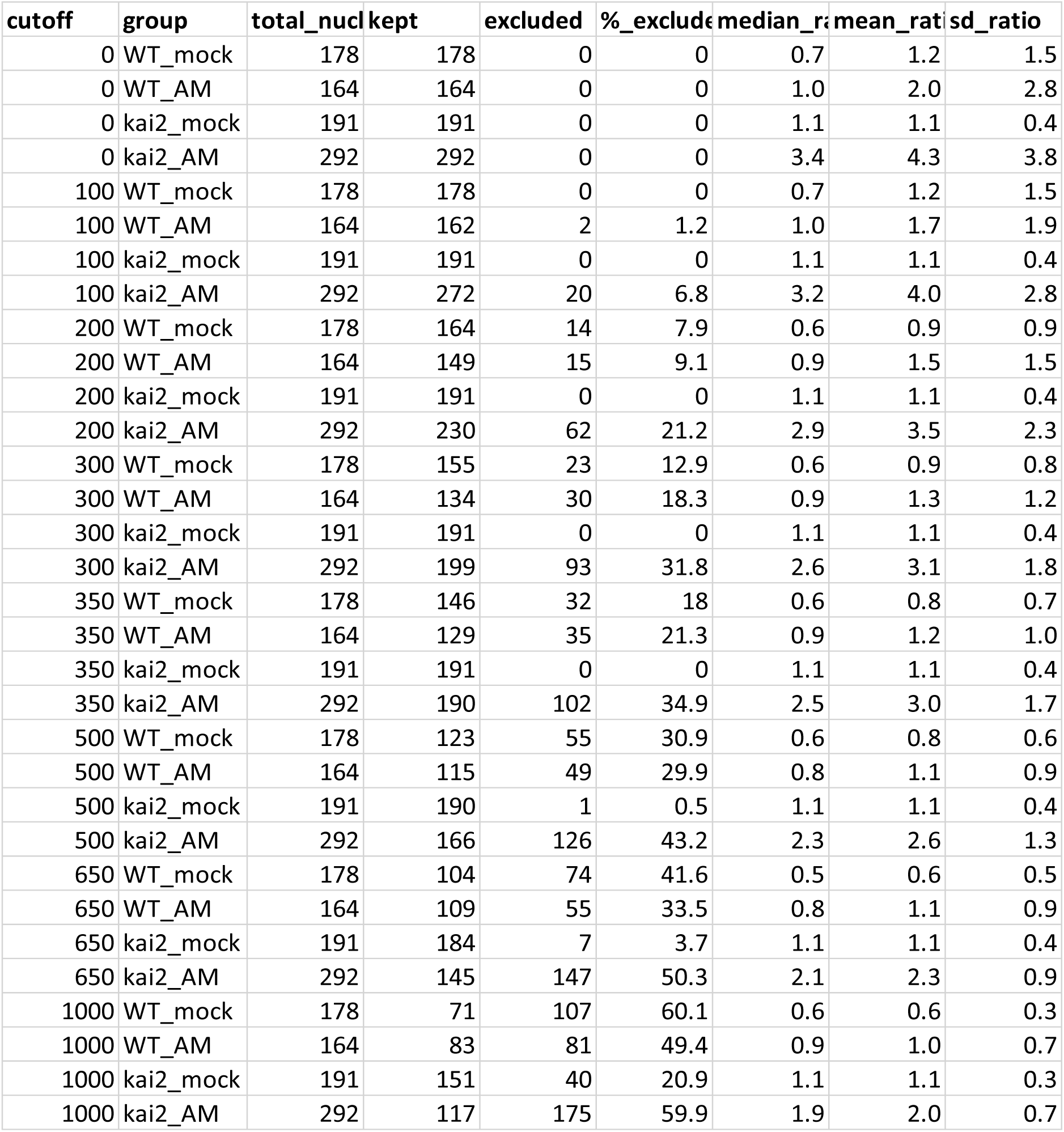
Number of excluded nuclei, median, mean and standard-deviation ratio at different minimum Venus cut-offs for analyses in Fig. 4.

**Supplementary Table 5.** Total number of DEGs in *L. japonicus* and *B. distachyon* transcriptome for *kai2* vs WT in mock conditions and total combined unique DEGs across all conditions.

| species | kai2_mock_vs_WT_mock | total |
| --- | --- | --- |
| *L. japonicus* | 2303 | 2537 |
| *B. distachyon* | 235 | 809 |

**Supplementary Table 6.** Distribution of nuclear mean intensity SMAX1_D2_-mScarlet-I to Venus ratio shown in Fig. 4c.

| group | n<1 | 1≤n≤2 | n>2 | %_<1 | %_1≤n≤2 | %_>2 |
| --- | --- | --- | --- | --- | --- | --- |
| WT_mock | 114 | 22 | 10 | 78.1 | 15.1 | 6.8 |
| WT_AM | 78 | 30 | 21 | 60.5 | 23.3 | 16.3 |
| kai2_mock | 50 | 136 | 5 | 26.2 | 71.2 | 2.6 |
| kai2_AM | 1 | 64 | 125 | 0.5 | 33.7 | 65.8 |

